# Super-resolution mapping of platelet α-granules uncovers non-polarized secretion and differential retention of VWF and thrombospondin-1

**DOI:** 10.64898/2026.10.16.759579

**Authors:** Smita Patil, Joran Deschamps, Massimiliano Garre, Shreya Sankar, Jonas Ries, Ingmar Schoen

**Author notes:** Correspondence: Ingmar Schoen, School of Pharmacy and Biomolecular Sciences, Irish Centre for Vascular Biology, Royal College of Surgeons in Ireland, 123 St. Stephen’s Green, Dublin 2, Ireland.

## Abstract

Platelet α-granules contain hundreds of bioactive proteins that regulate thrombosis, inflammation and tissue repair. The interplay between α-granule secretion and platelet biomechanics is central to platelet function but remains incompletely understood. Here we combined complementary super-resolution microscopy methods, computational image simulations and quantitative single-cell analysis to investigate α-granule nanoscale organization and secretion in tandem with cytoskeletal remodeling in spreading human platelets. We show that P-selectin, thrombospondin-1 (TSP1) and von Willebrand factor (VWF) occupy distinct subgranular domains and that apparent co-localization strongly depends on imaging resolution. We identify ring-like P-selectin staining as a robust signature of intact α-granules and demonstrate that α-granule secretion in spreading platelets was confined to the platelet cell body rather than the actively remodeling periphery. Quantitative assessment at the single-cell level showed that degranulation coincided with transitions from filopodia-rich to lamellipodial morphologies. Comparisons across adhesive substrates and perturbations by soluble agonists indicated that these processes are coordinated through overlapping signaling networks rather than direct local coupling. Analysis of secreted cargo revealed distinct extracellular fates despite no evidence for polarized secretion. TSP1 accumulated beneath platelets within diffusion barrier-restricted membrane pockets and around platelets following matrix binding, whereas VWF remained predominantly tethered to platelet membranes and formed short VWF strings. Together, our findings redefine the organization and release of platelet α-granules, demonstrate an absence of secretion polarity, and establish cargo-specific retention mechanisms that may influence platelet signaling and thrombus architecture.

**TEASER:** Super-resolution imaging uncovers non-polarized platelet α-granule secretion and cargo-specific extracellular deposition.

## INTRODUCTION

Platelets are anucleate blood cells and inherent multitaskers. At vascular lesions, platelets become activated to biomechanically form and stabilize the hemostatic plug, by binding to von Willebrand Factor (VWF) and fibrinogen, while secreting substances that modulate their own behavior as well as that of immune cells. Congenital platelet disorders exemplify the integral roles that biomechanics and exocytosis play in platelet function (*1, 2*). Platelet biomechanics is crucial for hemostasis (*3*), whereas platelet secretion can be friend or foe: it drives thrombosis and by (dis)engaging the immune system it can promote wound healing (*4, 5*), fight infections (*6, 7*) or facilitate cancer metastasis (*8, 9*). The exocytosis of α-granules, dense granules and lysosomes has been studied extensively in suspension in response to platelet agonists (*10–12*). This experimental approach investigates secretion in isolation, bereft of any influences that the extracellular surrounding and outside-in biomechanical signaling might have on secretion. Pioneering studies have started to investigate the spatiotemporal coordination of granule secretion within single spreading platelets (*13, 14*), however, fundamental questions on the relative timing and the spatial interdependence of biomechanical and secretory processes within single platelets remain unanswered, amongst others due to the difficulty to visualize specific components with sufficient spatial resolution.

The actin cytoskeleton plays a central role in biomechanical platelet responses, from initial platelet shape change and filopodia formation to the subsequent formation of lamellipodia and/or contractile stress fibers. Filamentous (f)-actin assembly and rearrangements are orchestrated by Rho family members, with Cdc42 mainly controlling filopodia, Rac1 stabilizing lamellipodia, and RhoA determining stress fiber formation and actomyosin activity (*15*). The actin cytoskeleton also plays a dual role in platelet exocytosis. On one hand, cortical actin is hindering the docking of granules to platelet membranes before Rac1 induces its breakdown by cofilin (*16*) and thus facilitates secretion (*17, 18*). On the other hand, actomyosin activity drives the initial centralization of α-granules which appears largely dispensable for α-granule secretion itself (*19–21*) while f-actin polymerization contributes to membrane fusion of α-granules (*22, 23*) but not dense granules (*24*). The dual role of f-actin explains a dose-dependent increase and then decrease of α-granule secretion at increasing concentrations of Latrunculin A (*25, 26*), a compound which perturbs f-actin assembly. Several components of the secretory machinery have been shown to interact with f-actin or f-actin binding proteins, including syntaxin-4 and VAMP-8 (*26*), syntaxin-11 and SNAP23 (*27*), syntaxin binding protein 5 (STXBP5) (*22*), and VAMP-7 (*13, 28*). Based on the peripheral localization of VAMP-7 (*29*) and defective lamellipodia formation in VAMP-7^-/-^ platelets, it has been proposed that ‘VAMP-7 links granule exocytosis with actin reorganization’ (*13*), while others argued that sequestration of VAMP-7 to the actin cytoskeleton might reduce its availability for granule secretion (*28*). Moreover, Rap1 is a main control switch of granule secretion in platelets but also prominently co-regulates integrin activation (*18, 30*). To what extent α-granule secretion on the one hand, and filopodia or lamellipodia formation on the other, influence each other through the shared f-actin pool and its interacting and regulatory proteins is thus incompletely understood.

The exocytosis of α-granules releases, amongst other cargos, adhesion proteins such as fibrinogen, fibronectin or VWF. When these adhesion proteins get immobilized at surfaces, they can mediate adhesion receptor binding and thus support (enable, accelerate, or increase) platelet adhesion and spreading, for example on uncoated glass (*31*). A ‘self-deposition’ of adhesion proteins enables single platelets to spread over non-adhesive areas in their immediate vicinity (*14*), while (re-)binding of adhesion proteins at the apical side of adherent platelets contributes to platelet accumulation at injury sites (*32, 33*). This raises the question whether platelets direct their secretion into specific directions once they adhere. An exclusive basolateral secretion from adherent platelets has been claimed (*14*) but was based on confocal microscopy which has very restricted spatial resolution in the z-direction. Moreover, other α-granule components can exert autocrine effects on platelets, much alike the secreted cargo from dense granules. Positive feedback loops include thrombospondin-1 (TSP1) dampening inhibitory cAMP signaling by binding to CD36 (*34*), soluble CD40 ligand (sCD40L) priming platelets through CD40-induced NF-κB signaling (*35*), or vascular endothelial growth factor (VEGF) boosting aggregation through Flt-1 and KDR receptor signaling on pre-activated platelets (*36*). Conversely, platelet-derived growth factor (PDGF) acts on PDGF α-receptors as a negative feedback signal (*37*). Since the effects depend on local concentrations of these molecules, it is important to better understand the timing and directionality of their release reactions which determine their effective accumulation.

Platelet α-granules are only 200-500 nm in diameter (*22*). They have been extensively studied using electron microscopy which provides nanometer resolution and cellular context, works well for suspended platelets, but lacks molecular specificity or suffers from poor (immunogold) labeling. Immunofluorescence and confocal microscopy offer molecular selectivity but are limited to ∼250 nm lateral and ∼800 nm axial resolution, making it difficult or impossible to assess the precise morphology of α-granules and the sub-localization of their contents, with two important consequences. One, in the absence of a morphological signature, any bright P-selectin staining might be mistaken as an α-granule, since P-selectin remains at the membrane after exocytosis. Two, in an anisotropic convoluted image, measurements of ‘co-localization’ (*38, 39*) are imperfect and ignore the inherently different localization of membrane proteins (such as P-selectin) and cargo proteins, of which some (such as VWF) only occupy sub-compartments within single granules (*10, 38, 40, 41*). More recently, super-resolution microscopy (SRM) methods and expansion microscopy (ExM) have been adopted to study the platelet cytoskeleton (*42–47*) or granules (*48–52*). The ∼2-fold resolution improvement of structured illumination microscopy (SIM) led to the discovery of actin-nodules in platelets (*45*), confirmed the excentric localization of VWF and VWF-propeptide in α-granules and their respective release kinetics (*52*), and emphasized the mesh-like morphology of P-selectin on membranes of close-by α-granules (*49, 50*), which is also seen to some extent in good-quality confocal images (*28*). SRM techniques which routinely reach a resolution better than 100 nm have elucidated extracellular matrix deposition by platelets (*33*), cytoskeletal and organelle arrangements (*42, 46*) and cell surface receptor clustering (*53*) in spreading platelets, as well as receptor and organelle organization in suspended platelets (*44, 47, 54*). The full potential of SRM to elucidate the interrelationship between platelet biomechanics and secretion has so far remained underexploited.

In this study, we use a combination of different state-of-the-art SRM methods, namely lifetime (tau) STimulated Emission Depletion (τSTED) and direct STochastic Optical Reconstruction Microscopy (dSTORM), in combination with advanced image analysis and computer simulations to revisit α-granule cargo arrangement in resting platelets as well as α-granule secretion and actin cytoskeletal rearrangements in spreading platelets. Our improved image quality and quantitative evaluation reveal new insights into the spatiotemporal coordination of these processes in single platelets and help to resolve existing ambiguities.

## RESULTS

### τSTED microscopy resolves distinct morphologies of α-granule components in resting platelets

We first sought to characterize α-granule components in intact α-granules by 2D τSTED microscopy. We chose P-selectin as a prototypical α-granule membrane protein and TSP1 as an abundant cargo protein. In resting platelets, P-selectin immunostaining appeared as a web-like mesh composed of closely packed rings (Figure 1A). Rings were not stained homogeneously but frequently had gaps in the staining. These rings were relatively round, had a length of 318±71 nm (mean±std) and an apparent wall thickness of 50–100 nm (Figure 1B). This wall thickness was in the range of the optical resolution of 2D τSTED images according to point spread function (PSF) measurements and the Fourier ring correlation (FRC) resolution metric (Supplementary Figure 1), meaning that the underlying structures were actually thinner. These P-selectin images and their interpretation agree with previous SRM studies (*28, 49, 50*). Immunostaining of the cargo protein TSP1 showed spots of heterogeneous size, shape, intensity and texture, with larger spots being brighter and more homogeneous (Figure 1C). Spots were 309±85 nm long on average, mostly roundish although some were more elongated, and were often packed closely (Figure 1D), which would render them indistinguishable at lower optical resolution.

**Figure 1.**
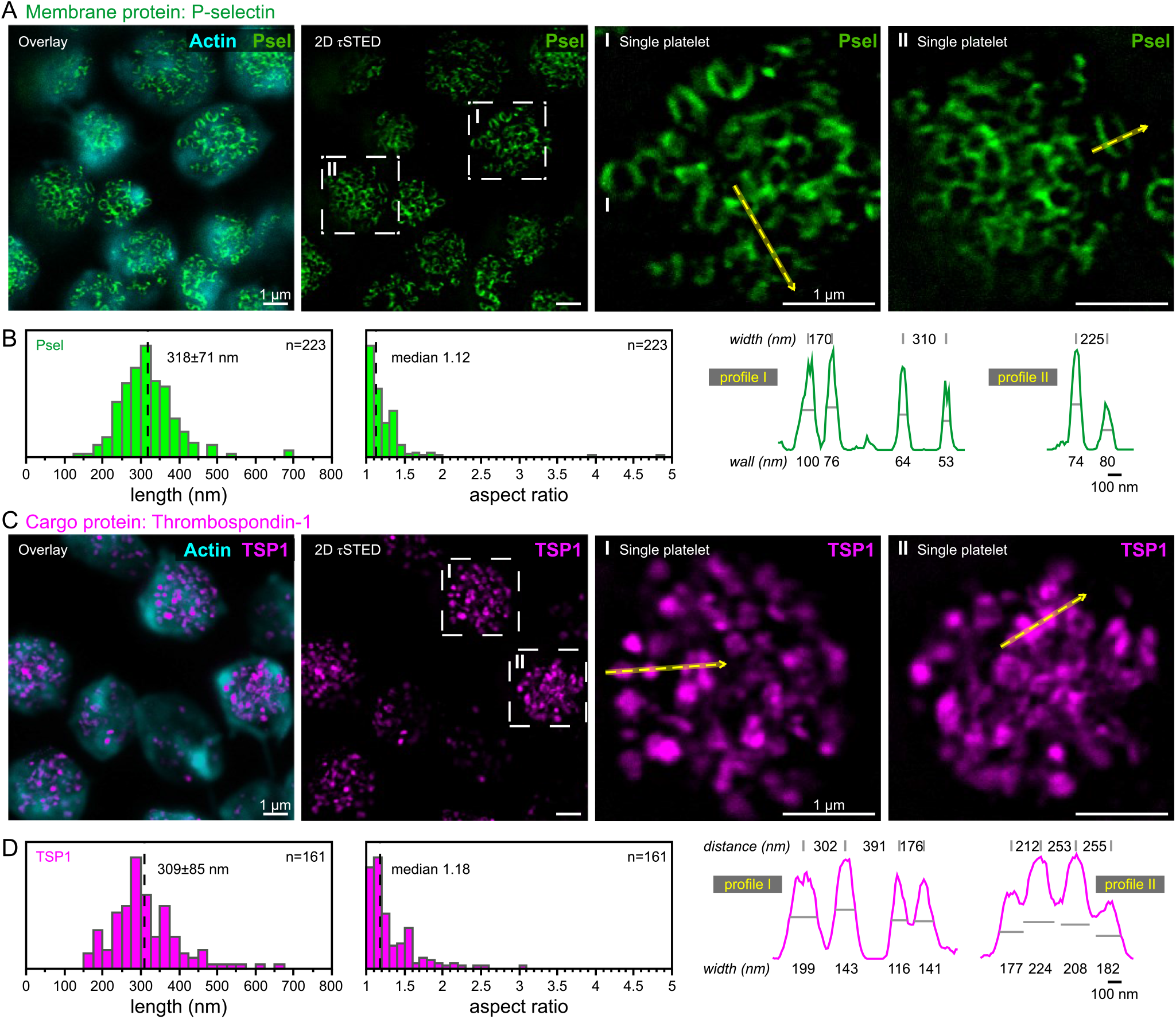
τSTED microscopy resolves distinct morphologies of α-granule components in resting platelets. Washed human platelets were fixed in solution, immobilised onto coverslips, permeabilized, immunostained, a d imaged by confocal (f-actin; cyan) or 2D τSTED microscopy (granule components). (**A**) Representative overview and magnified single cell images of P-selectin staining (Psel; green). Closely packed ring-like staining is observed. Scale bars: 1 µm. (**B**) Dimensional analysis of P-selectin rings. Left: length and aspect ratio. Right: intensity line profiles along dashed yellow arrows in (A). P-selectin rings had diameters of 200–400 nm, their wall thickness was in the range of the optical resolution of 2D τSTED. Scale bar: 100 nm. (**C**) Representative overview and magnified single cell images of thrombospondin-1 (TSP1; magenta). Spotty punctuated staining is observed. Scale bars: 1 µm. (**D**) Dimensional analysis of thrombospondin-1 puncta. Left: length and aspect ratio. Right: intensity line profiles along dashed yellow arrows in (C). Thrombospondin-1 puncta had diameters of 150–350 nm and could appear in close proximity to each other. Scale bar: 100 nm.

To further rationalize the appearance of α-granule components in 2D τSTED images, we next performed Monte Carlo computer simulations to generate realistic granule images from first principles (Figure 2A). We modeled granules as spheres, distributed α-granule proteins proportionally between granules according to their size distribution per platelet (*22*), and randomly scattered them over the surface (membrane proteins) or throughout the inner volume (cargo proteins). Taking into account the finite labeling efficiency of antibody-epitope binding and a variable number of fluorophores per antibody, the ground truth 3D distribution of fluorescent emitters was computed and convolved by the experimental PSF. Finally, expected photon counts per pixel were exposed to Poisson noise to capture the stochastic photon emission process. Simulated images of P-selectin showed characteristic rings (Figure 2B) which were the result of an effective z-projection due to the axially elongated PSF of 2D τSTED. Qualitative comparison of simulated images with their experimental counterparts (see Figure 1A) suggested a relatively high labeling efficiency for P-selectin in our experiments. Even at these high labeling efficiencies, the limited number of P-selectin molecules per granule and their random positions led to staining irregularities along the ring, but rarely to larger gaps as observed in Figure 1A. Measurements of the ring diameter systematically underestimated the ground truth granule size by ∼50 nm, as expected from the projected spherical geometry and the PSF convolution. Simulated images of TSP1 (Figure 2C) showed round spots throughout. Spots had an increasingly irregular texture with decreasing labeling efficiency, but not as spotty as for P-selectin due to the much higher protein copy numbers per granule. A qualitative comparison to experimental images (see Figure 1C) suggested an intermediate immunolabeling efficiency for TSP1 in our experiments. Measurements of the full-width-at-half-maximum (FWHM) of TSP1 spots systematically underestimated the ground truth granule size by ∼70 nm. Defocusing mainly affected the brightness but only marginally the ring-like appearance of P-selectin or the spot-like appearance of TSP1 (Supplementary Figure S2A).

**Figure 2.**
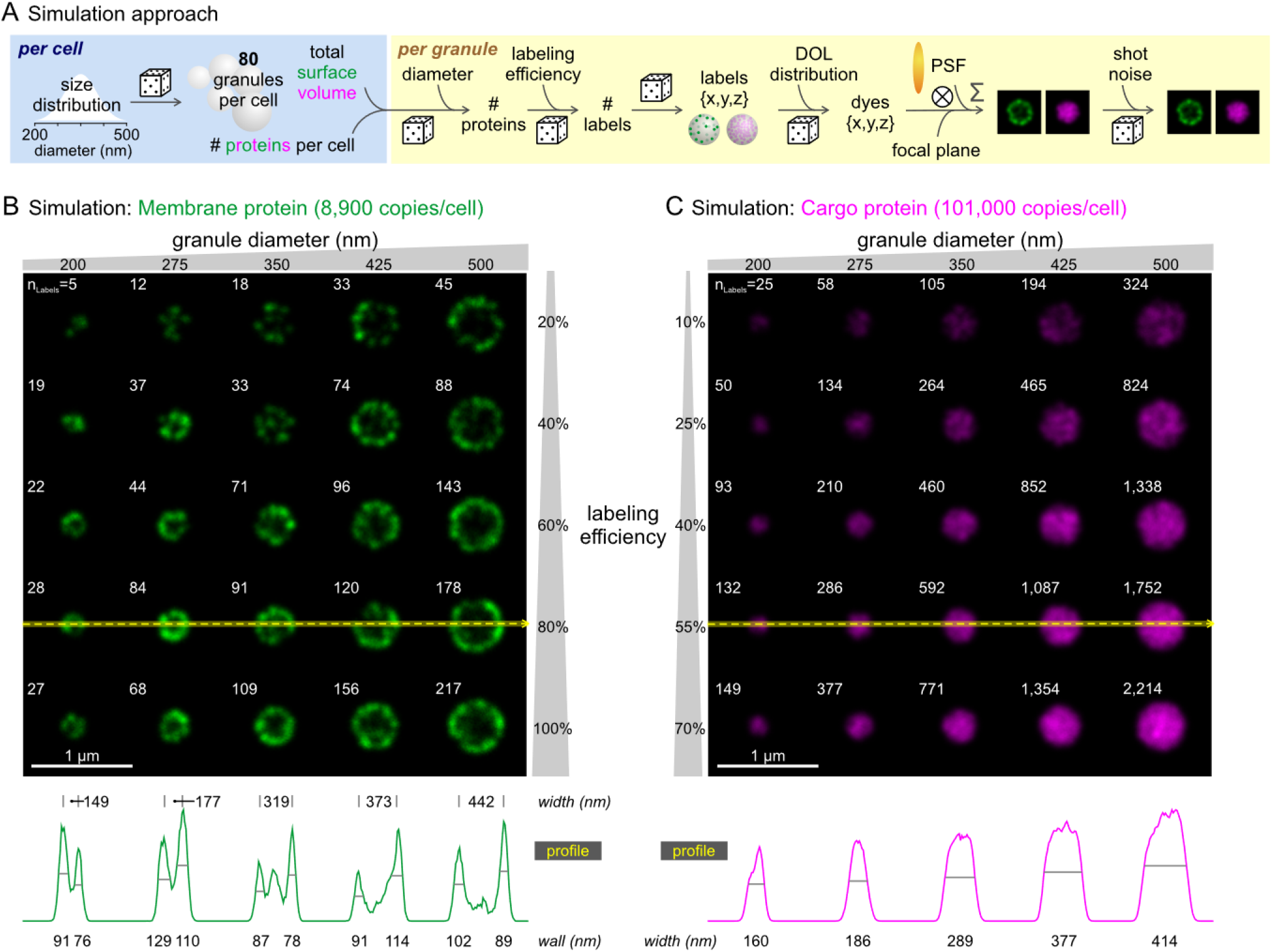
Monte Carlo simulations reproduce key features of experimental α-granule images. (**A**) Simulation approach. α-granules were simulated as hollow (membrane protein) or solid (cargo proteins) spheres with randomly sampled size, labeling efficiency, label distribution, and fluorophore number. This 3D distribution of fluorescent point sources was then convolved by the measured 3D point-spread function of 2D τSTED relative to the focal plane, and photon shot noise was added. For further details, see the methods section. (**B**) Simulated images of a membrane protein for different α-granule diameters and immunolabelling efficiencies. The protein copy number per cell was based on proteomic data for P-selectin in human platelets. The membrane protein staining appears as a ring; the limited label number and their random distribution can compromise the smooth ring-like appearance. Dimensional analysis of P-selectin rings using the intensity line profile along dashed yellow arrow confirms wall thicknesses in the range of the optical resolution of 2D τSTED, and diameters that slightly underestimate the true granule size. Scale bar: 1 µm. (**C**) Simulated images of a cargo protein for different α-granule diameters and immunolabelling efficiencies. The protein copy number per cell was based on proteomic data for thrombospondin-1 in human platelets. The cargo protein staining appears as a spot; a very low labelling efficiency can compromise the smooth appearance. Dimensional analysis of thrombospondin-1 spots using the intensity line profile along dashed yellow arrow results in spot diameters that slightly underestimate the true granule size, similar to the analysis in (B). Scale bar: 1 µm.

Based on the very good agreement of experimental and simulated images of P-selectin, we conclude that the ring-like morphology of P-selectin in 2D τSTED images is an essential hallmark of intact granules which can help to distinguish them from non-granular P-selectin staining. The limited agreement on the shape of TSP1 spots suggests that TSP1 might not occupy the whole interior of α-granules.

### In intact α-granules, VWF localizes separately from P-selectin and TSP1

In electron microscopy images, VWF occupies a densely packed peripheral sub-region within a subset of α-granules (*38, 40, 41*). 2D τSTED images of resting platelets showed elongated and often curved VWF staining (Figure 3A) of varying length (100–400 nm) and width (80– 150 nm) (Figure 3B). To visualize the relative location of VWF within granules, we performed dual color 2D τSTED of VWF and P-selectin (Figure 3C). While few VWF staining was apparently located outside of granules (’o’), most VWF staining localized close to one (asterisks) or more (’∞’) sides of individual P-selectin rings, consistent with its known localization to peripheral sub-regions (*52*). Strikingly, P-selectin staining was often absent or reduced proximal to VWF (arrowheads). This systematic observation indicates that gaps in the P-selectin staining might reflect an uneven distribution of P-selectin over the granule surface rather than stochastic fluctuations (see Figure 2B), which could be related to the packaging of cargo within granules.

**Figure 3.**
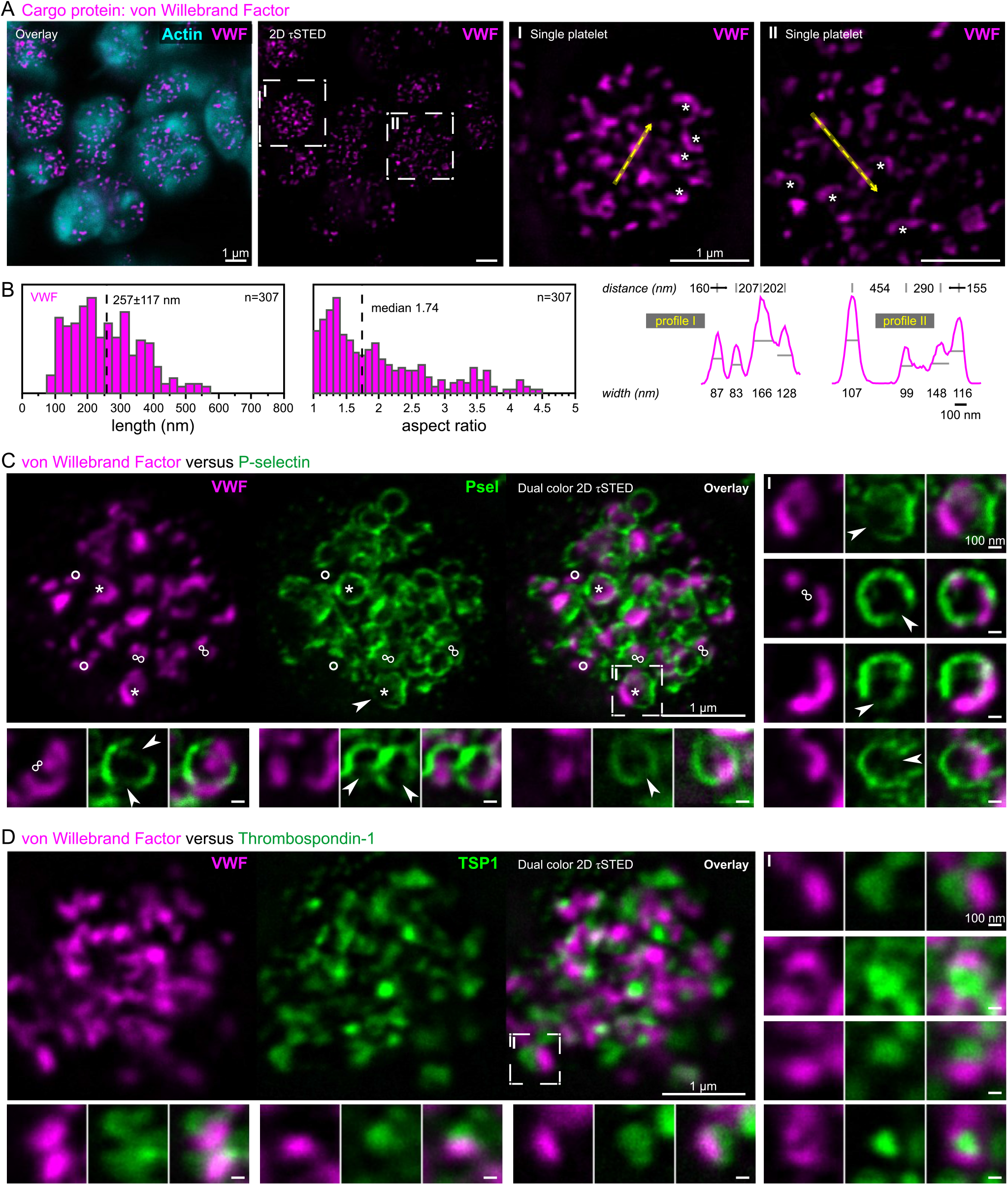
In intact α-granules, VWF localizes separately from P-selectin and TSP1. Washed human platelets were fixed in solution, immobilised onto coverslips, permeabilized, immunostained, and imaged by confocal (f-actin; cyan) or 2D τSTED microscopy (granule components). (**A**) Representative overview and magnified single cell images of VWF staining (magenta). Separate, thin, elongated, often croissant-shaped (asterisks) staining is observed. Scale bars: 1 µm. (**B**) Dimensional analysis of VWF spots. Left: length and aspect ratio. Right: intensity line profiles along dashed yellow arrows in (A). VWF spots were up to 400 nm long and 80–150 nm wide. Scale bar: 100 nm. (**C**) Representative single cell and magnified single granule dual-color 2D τSTED images of VWF (magenta) and P-selectin (green). VWF localized peripherally within single granules (asterisk). P-selectin staining was often reduced at VWF locations (arrowheads). Some granules contained more than one VWF spot (‘∞‘). Some VWF had no corresponding P-selectin staining (‘o‘). Scale bars: 1 µm (cell), 100 nm (insets). (**D**) Representative single cell and magnified single granule dual-color 2D τSTED images of VWF (magenta) and thrombospondin-1 (TSP1; green). VWF localized in close proximity to TSP1, potentially implying co-occupation of same granules, but did not overlap, implying distinct storage sub-compartments. Scale bars: 1 µm (cell), 100 nm (insets).

VWF also in relation to TSP1 occupied distinct locations within the platelet and proposedly within single granules (Figure 3D), however, limitations in antibody orthogonality and the number of STED channels did not allow us to image all three stains in the same sample. We conclude that at the superior resolution of 2D τSTED, no true co-localization (overlap) is observed between VWF and TSP1, whereas co-occupation (sharing) of the same granules is plausible.

### Spreading platelets secrete α-granules exclusively from their cell body

To investigate the temporal relationship between α-granule secretion and actin dynamics during spreading, we seeded washed human platelets for 2–20 minutes on fibrinogen-coated coverslips and imaged the fixed, permeabilized and stained samples by dual-color 2D τSTED (Figure 4A). Cells spread for only 2 minutes were either small with filopodia or had already started to spread. The P-selectin staining in spreading platelets varied greatly from cell to cell. In some spread cells, the characteristic ring-shaped P-selectin mesh was observed, very much like in resting platelets (see Figure 1A), indicating largely intact α-granules. This granulomere region (Figure 4A, dashed outline) was usually surrounded by branched actin or stronger actin bundles, with only few traversing actin filaments. At later time points, nearly all platelets were fully spread, but the P-selectin staining remained heterogeneous. In many cells, the central strong P-selectin staining had completely disappeared, indicating all α-granules had been secreted, whereas in other cells, few intact P-selectin rings remained. Nearly all cells at this late time point exhibited a weak disperse spotty P-selectin staining throughout the lamellipodium (asterisks) and on some filopodia (arrowheads). We note that this weak staining was specific for P-selectin and not a result of bleed-through or unspecific background, which were corrected for or negligible, respectively (Supplementary Figures S3+S4).

**Figure 4.**
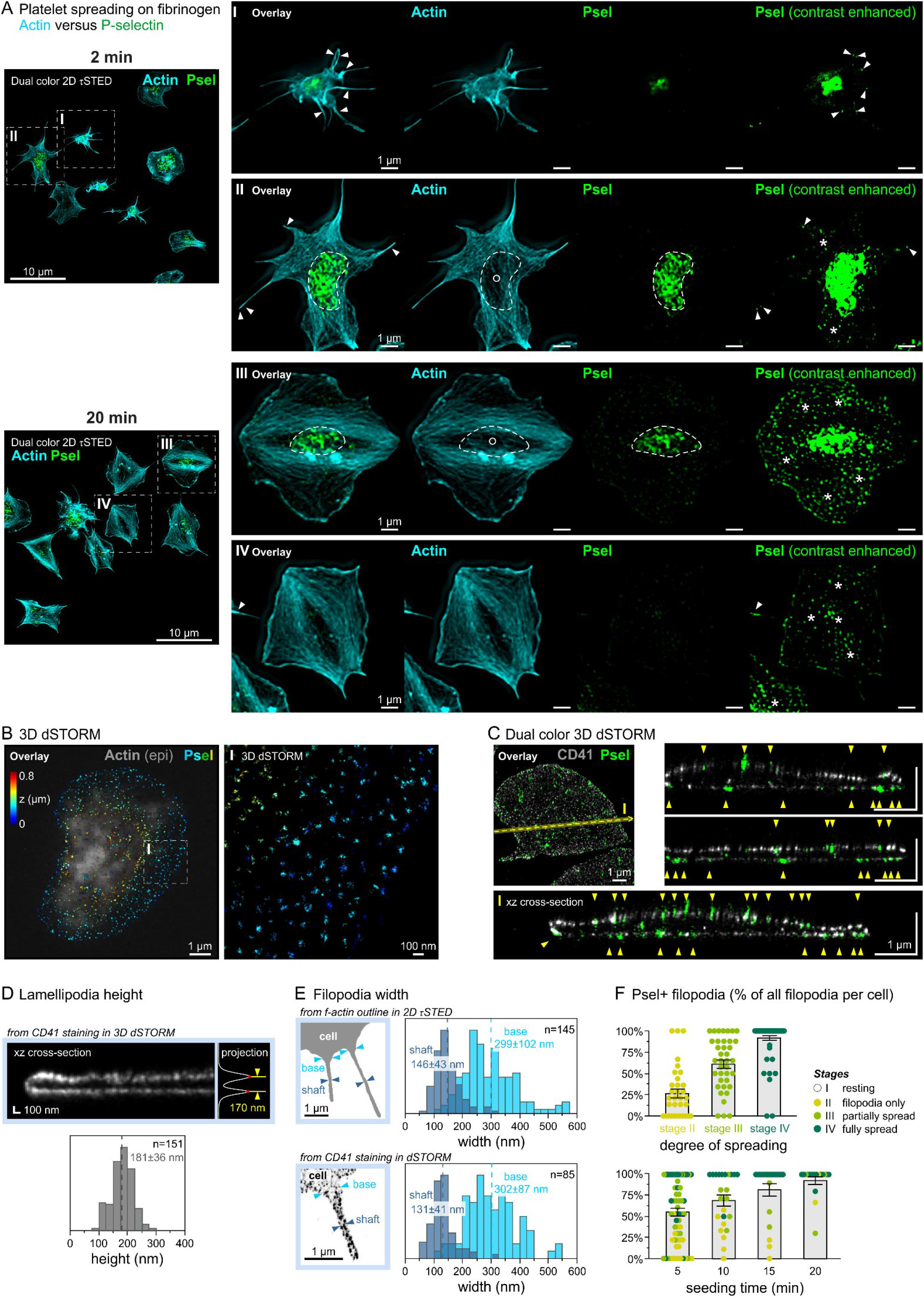
Spreading platelets secrete α-granules exclusively from their cell body. Washed human platelets were spread on fibrinogen-coated coverslips for up to 20 min, fixed at different time points, permeabilized, immunostained, and imaged by SRM. (**A**) Representative overview images and magnified single cell insets of dual-color 2D τSTED f-actin (cyan) and P-selectin (Psel; green) images after spreading for 2 or 20 minutes. Contrast-enhanced P-selectin images (right) show occasional punctate P-selectin staining along filopodia (arrowheads), an increase in disperse staining (asterisks) across the whole cell, and a decrease of intact granules at the cell centre. F-actin was largely excluded from the granulomer region (‘o‘). (**B**) Representative single cell 3D dSTORM image of P-selectin (color-coded according to z-position) after 20 min. The overlay shows f-actin (gray) for context. The high sensitivity and resolution of dSTORM confirms the disperse character of P-selectin staining. (**C**) Single cell dual color 3D dSTORM image of P-selectin (green) and integrin α_IIb_ (CD41; gray) and xz cross-sectional sideviews. The superior z-resolution of dSTORM confirms co-localization of the disperse P-selectin staining with the plasma membrane (yellow arrowheads). (**D**) Determination of lamellipodia height from 3D dSTORM xz cross-sections of integrin α_IIb_ (CD41; gray). The height ranged from 100–250 nm. (**E**) Determination of filopodia width from 2D τSTED images of f-actin (top) or dSTORM images of integrin α_IIb_ (CD41; bottom). The shaft was up to 200 nm wide, the base up to 400 nm. (**F**) Percentage of filopodia (per cell) that were decorated with P-selectin, categorised by spreading stage (top, color-coded) and spreading time (bottom). The decoration of filopodia with P-selectin was low at initial stages and high in fully spread platelets.

Based on the distinct appearance in terms of signal strength and morphology, we attributed this disperse weak P-selectin staining to membrane-expressed P-selectin. To confirm this interpretation, we used another SRM method, 3D dSTORM, which has single molecule sensitivity and reaches a resolution of 30–40 nm laterally (Supplementary Figure S5) and better than 100 nm axially. The higher sensitivity revealed a dense coverage of the lamellipodium by the disperse P-selectin staining (Figure 4B), while the superior z-resolution and a counter-staining for platelet integrins clearly demonstrated that this P-selectin staining localized to both the basolateral and apical membranes of lamellipodia and of the cell body (Figure 4C). Since unspecific staining was negligible (Supplementary Figure S3B), these data confirmed that the disperse P-selectin staining resembled membrane-expressed P-selectin after α-granule exocytosis.

A previous study proposed that sites of α-granule secretion could template the formation of filopodia or lamellipodia, facilitated by the dual role of VAMP-7 in membrane fusion and cytoskeletal rearrangements (*13*). Since VAMP-7 and P-selectin staining were detected in the lamellipodia of spread platelets, it was suggested that a subpopulation of α-granules could be transported there (*29*). A systematic measurement of the lamellipodium height using the membrane staining in our 3D dSTORM data yielded an average height of 181±36 nm (Figure 4D), confirming previous measurements (191±37 nm) based on lamellipodial f-actin (*33*). Considering the larger size of intact α-granules (200–400 nm, see Figure 1B), and consistent with the total absence of P-selectin rings in lamellipodia (see Figure 4A), it is questionable whether α-granules can be transported through the lamellipodium without inducing their fusion with the membrane. The measured width of filopodia shafts was mostly between 75–200 nm, in excellent agreement with electron microscopy data (*55*), whereas their base was 180–400 nm wide (Figure 4D). Consistent with the larger size of α-granules, we never observed full P-selectin rings within the filopodia shaft, but also never at their base, including in unspread platelets. We also never observed an enrichment of P-selectin along a filopodium as it could be expected if a filopodium was formed in short succession after a fusion event from the locally externalized P-selectin-positive α-granule membrane. The disperse P-selectin staining observed along filopodia was largely absent on newly formed filopodia at early spreading stages but increased over time (Figure 4E), consistent with membrane-expressed mobile P-selectin diffusing into filopodia over time.

Taken together, α-granule exocytosis exclusively occurred from the cell body in unspread platelets or from the granulomere in spread platelets, but not from peripheral f-actin-filled lamellipodia or filopodia.

### α-granule secretion correlates temporally with filopodia and lamellipodia formation

The spatial incompatibility of thin filopodia and lamellipodia with α-granules does not preclude a coordination between cytoskeletal rearrangements and degranulation through other mechanisms. α_IIb_β_3_ integrin outside-in signaling (*56*), mechanosensing (*14, 57, 58*), and secondary mediators (*12*) initiate and orchestrate filo-/lamellipodia formation and granule secretion in platelets spreading on fibrinogen, whereas on collagen, α_2_β_1_ integrins mediate spreading while GPVI signaling provides additional stimuli (*53, 59*). We thus next sought to characterize the dynamics of cytoskeleton reorganization and degranulation on these two adhesion protein coatings in order to identify overarching interdependencies.

Taking advantage of our 2D τSTED image resolution, we implemented an automated detection of filo-/pseudopodia (Figure 5A; see Methods) for platelets spreading on fibrinogen- and collagen-coated glass, extending our abilities to analyze platelet morphology in a high-throughput fashion (*46*). The number of filopodia per platelet varied greatly among platelets fixed at the same time point. The average number of filopodia per platelet decreased over time on either adhesion protein, but platelets on collagen initially formed ∼2 filopodia more on average (Figure 5B) and filopodia were consistently longer by ∼0.5 µm (Figure 5C). These differences in filopodia formation were also reflected by a decreased circularity on collagen and an initially decreased f-actin alignment (Supplementary Figure S6 A+B). To estimate the degree of α-granule secretion in spreading platelets, we took advantage of our key finding that P-selectin distributes evenly across spread platelets after secretion whereas intact granules are exclusively located in the central granulomere region (see Figure 4). The ratio of the P-selectin intensity in peripheral regions of the cell divided by the intensity of the inner (central) region showed very good agreement with a visual assessment of microscopy images, with ratios <0.5 indicating largely intact granules, and values approaching ∼1 reflecting complete secretion (Figure 5D). This peripheral/inner P-selectin ratio of platelets varied greatly at specific time points, reflecting the observed secretion heterogeneity (see Figure 4A). On fibrinogen, the average peripheral/inner P-selectin ratio increased from ∼0.5 at 2 minutes to ∼0.8 at 15 minutes and then remained steady (Figure 5E). On collagen, more platelets had undergone secretion already at 2 minutes (∼0.8 on average) and all platelets had secreted (∼1.0) after 10 minutes. Adhesion protein identity thus affected both, filopodia formation and granule secretion.

**Figure 5.**
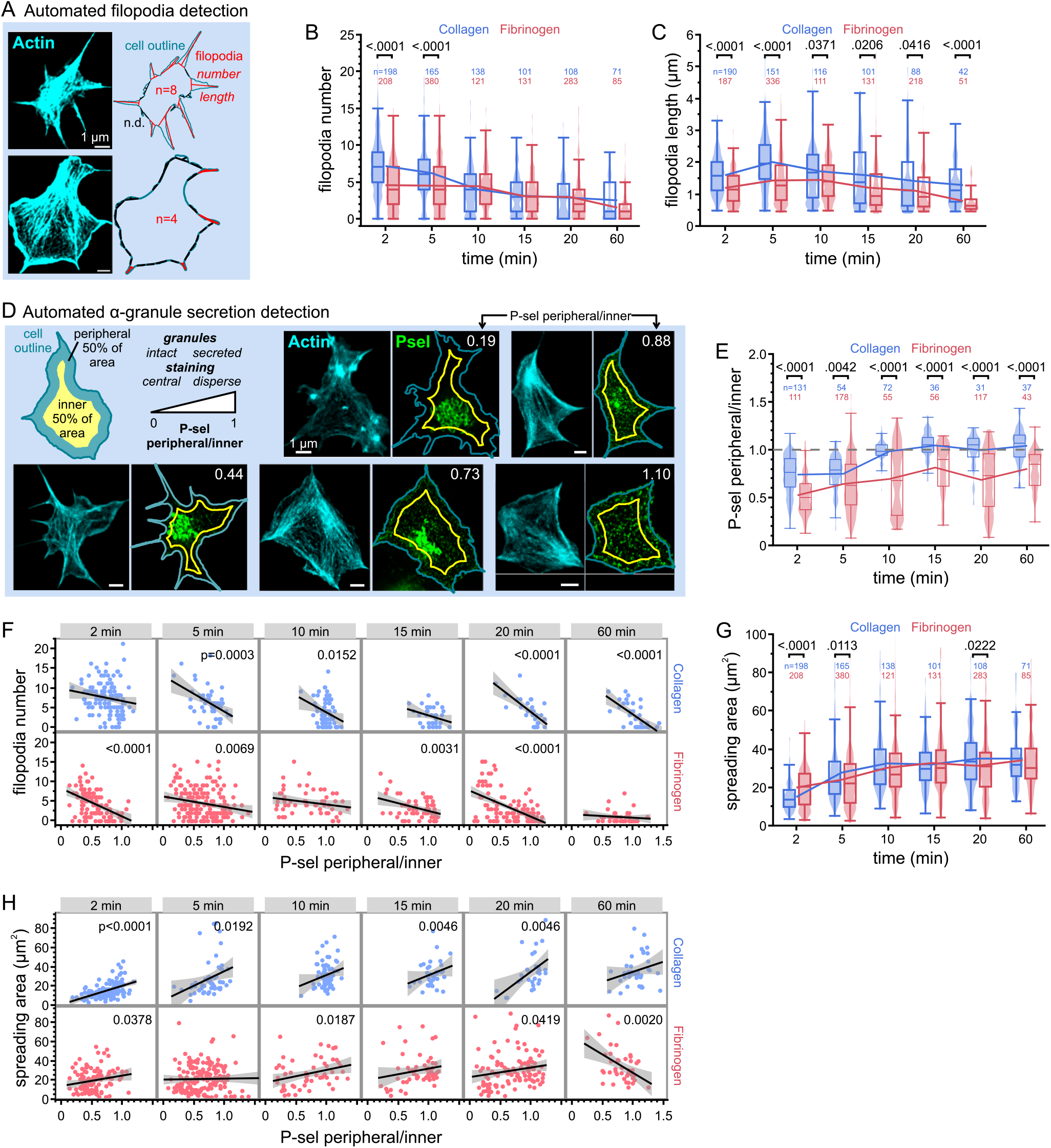
α-granule secretion correlates temporally with filopodia and lamellipodia formation. Washed human platelets were spread on fibrinogen- or collagen I-coated coverslips up to 60 min, fixed at different time points, permeabilized, immunostained, and imaged by 2D τSTED (**A**) Illustration of the automated filopodia detection from cell outlines of f-actin 2D τSTED images (**B**) Number and (**C**) mean length of filopodia per cell for platelets spread on fibrinogen (red) or collagen I (blue). Single cell data are shown as violin plots and summarised as box plots (25/50/75 percentile, whiskers excluding outliers). Connecting lines depict the mean per time point. The total number of platelets per time point pooled from 1-3 independent experiments is denoted above the graphs. Statistical comparisons between adhesion proteins were performed by student’s t-tests. Platelets on collagen I initially formed more filopodia than on fibrinogen. The number of filopodia decreased over time on both adhesion proteins. Filopodia were consistently longer on collagen I. (**D**) Illustration of the automated detection of α-granule secretion from cell outlines and P-selectin 2D τSTED images. Example images show the correlation of the P-sel peripheral/inner metric with the P-selectin distribution within the cell: low values indicate an intact granulomer, values around 1 complete secretion. (**E**) P-selectin peripheral/inner ratio for platelets spread on fibrinogen (red) or collagen I (blue). Data visualisation and statistical comparisons as in (**B**). The total number of platelets per time point pooled from 1-3 independent experiments is denoted above the graphs. The α-granule secretion proceeded over the spreading duration on either adhesion coating, but secretion in platelets on collagen I was faster and more complete. (**F**) Correlation analysis of filopodia number versus the degree of α-granule secretion (measured by the peripheral-to-inner P-selectin intensity ratio). Each data point represents a single cell; total cell numbers as in (E). Linear regression (black line) and 95% confidence intervals (gray) show an anti-correlation for all time points and on either adhesion protein coating, platelets with less filopodia showed more complete secretion. (**G**) Platelet spreading area. Data visualisation and statistical comparisons as in (B). Spreading was comparable on fibrinogen and collagen I. (**H**) Correlation analysis of spreading area versus the degree of α-granule secretion (measured by the peripheral-to-inner P-selectin intensity ratio). Visualisation as in (F). A positive correlation was consistently observed, with more spread platelets showing complete secretion.

We next asked how cytoskeletal reorganization and degranulation were interrelated on the single platelet level. Taking advantage of the observed platelet population heterogeneity and the simultaneous measurement of morphology and degranulation within the same platelet, we performed a correlation analysis between the filopodia number and the P-selectin peripheral/inner ratio per time point (Figure 5F). Platelets with smaller P-selectin ratios had on average more filopodia than platelets that had fully secreted. This correlation was consistently observed across time points on collagen and fibrinogen. As a proxy of lamellipodia formation, we analyzed the spreading area of individual cells. Platelets spread mainly during the first 10 minutes, with a head start on fibrinogen, but reaching very similar final spread areas (Figure 5G). The spreading area correlated positively with the degree of α-granule secretion on a single platelet level (Figure 5H). Finally, the number of filopodia was lower in more fully spread platelets (Supplementary Figure S6C).

In summary, α-granule secretion within single spreading platelets coincided and correlated with filopodia number and lamellipodia formation, both temporally and within the cell population. However, these correlations did not strictly hold when comparing the results across different adhesion proteins: while initial degranulation activity and final degranulation level were higher and more complete on collagen compared to fibrinogen, the initial filopodia number was also higher but the final spread area was the same.

### Secreted TSP1 accumulates beneath and around spread platelets

TSP1 modulates platelet reactivity (*34*), endothelial cell angiogenesis (*60*), and immune cell activation (*61*). TSP1 binds to the extracellular matrix (ECM) (*62, 63*) and exerts signaling effects through CD36 on platelets (*64*) but its fate after α-granule secretion from spreading platelets and potential autocrine effects remain largely unclear. Unspread platelets on fibrinogen at early time points after seeding showed granular TSP1 staining in their cell body (Figure 6A), qualitatively similar to TSP1 staining in resting platelets (see Figures 1+3). With increasing time during platelet spreading, the central TSP1 staining appeared more pronounced and fused, with prominent ‘pockets’ that exceeded 1 µm laterally (Figure 6A, arrowheads) and that were located between f-actin bundles (dashed outlines). Most lamellipodia were completely devoid of TSP1 (‘o’), while TSP1 accumulated at the fibrinogen coating around spread platelets over time (asterisks). The observed gradient of decreasing TSP1 intensity away from the cells is consistent with TSP1 being secreted by these cells and diffusing before binding to the adhesion protein coating.

**Figure 6.**
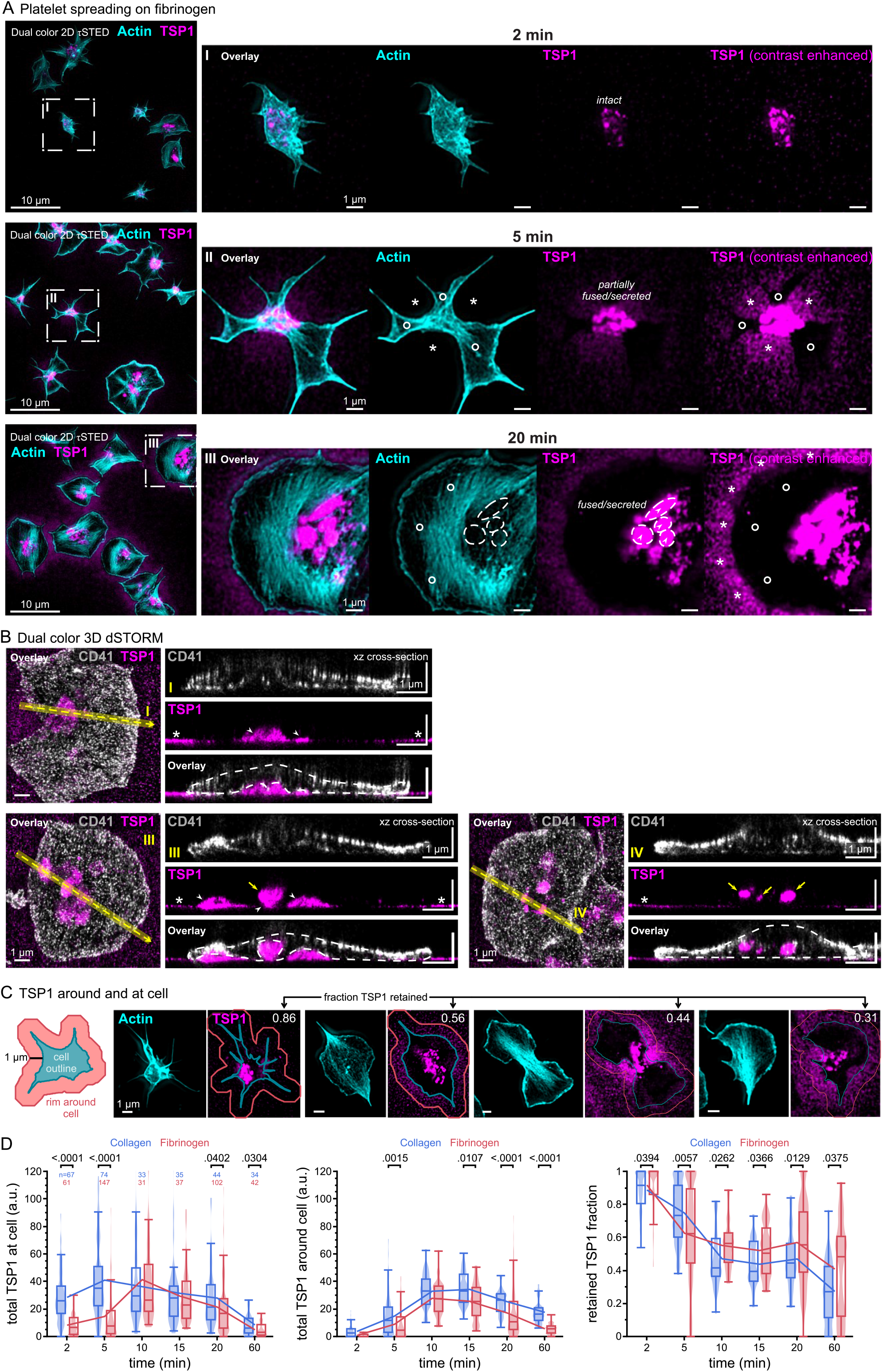
Secreted TSP1 accumulates beneath and around spread platelets. Washed human platelets were spread on fibrinogen- or collagen I-coated coverslips up to 60 min, fixed at different time points, permeabilized, immunostained, and imaged by SRM. **A**. Representative overview images and magnified single cell insets of dual-color 2D τSTED f-actin (cyan) and TSP1 (magenta) images after spreading for 2, 5 or 20 min on fibrinogen. Contrast-enhanced TSP1 images (right) show disperse TSP1 staining (asterisks) accumulating around spread platelets over time, whereas regions below lamellipodia are largely devoid of TSP1 (‘o‘). Bright staining is retained at the cell centre. **B**. Single cell dual color 3D dSTORM image of TSP1 (magenta) and integrin α_IIb_ (CD41; gray) and xz cross-sectional sideviews for platelets spread on fibrinogen. The superior z-resolution of dSTORM distinguishes between TSP1 accumulating at the coverslip (asterisks), intracellular granules (yellow arrows), pockets of TSP1 being retained beneath the basal plasma membrane (arrowheads), and occasional fusion channels between granular and secreted TSP1 (‘o‘). **C**. Illustration of the estimate of TSP1 retention at (within or below) the cell, as compared to its accumulation around the cell in a 1 µm wide rim. Example images show the correlation of the ‘retained TSP1 fraction‘ metric with the visual appearance / distribution of TSP1: low values indicate cells largely devoid of TSP1. **D**. Total TSP1 intensity at (left) or around (middle) the cell and their ratio (right). TSP1 was secreted faster initially at the cell and accummulated more on collagen I compared to fibrinogen. Accordingly, more TSP1 was retained within and beneath platelets on fibrinogen.

Given that most platelets degranulate completely after 20 minutes (see Figures 4A+5E), we further investigated the nature of the central persistent TSP1 staining using 3D dSTORM with a CD41 membrane counterstain (Figure 6B and Supplementary Figure S7). Cross-sectional sideviews demonstrated that in many instances, the fused TSP1 staining was contained within ‘pockets’ (arrowheads) formed over the coverslip coating by the upward curved basal membrane, sealed off by the normally very close contact between basal membrane and coverslip coating. Because of the high z-resolution, these pockets were clearly distinguished from intact granules within the cell (yellow arrows). In one instance, they were connected to each other by an apparent fusion channel (white arrow). Beside these pockets, TSP1 was rarely seen at the coverslip beneath lamellipodia but always around cells (asterisks). TSP1 nearly never directly localized to the platelet membrane.

To quantify the retention of TSP1 within and beneath spreading platelets, we integrated the TSP1 fluorescence intensity within the cell outline and divided it by the integrated fluorescence intensity of a larger region comprising the cell’s immediate surrounding. This TSP1 intensity ratio correlated qualitatively with the visual impression of TSP1 retention within/beneath the cell, with ratios >0.8 indicating largely cellular localization, and values <0.5 signifying substantial release of TSP1 into the surrounding (Figure 6C). In platelets on fibrinogen, the retained TSP1 fraction dropped over the first 10 minutes, then plateaued, before reducing further towards 60 minutes (Figure 6D). This observation reflected the degranulation activity over the first 10 minutes and a very slow lateral diffusion of basally secreted TSP1. Platelets on collagen followed a similar trend yet reached a lower plateau and a lower final retention (Figure 6D), in agreement with a more complete secretion (see Figure 5E).

In summary, our data show that platelets secreted TSP1 into all directions but with different outcomes: apically (‘upwards’) or laterally (‘sideways’) secreted TSP1 diffused away and bound to the ECM, while basally (‘downwards’) secreted TSP1 was retained for prolonged times in membrane pockets beneath the platelet. A direct rebinding of TSP1 to platelet membranes was rarely observed.

### Secreted VWF is retained at apical and basal platelet membranes

VWF secretion from endothelial cells is polarized: upon stimulation, high molecular weight VWF from Weibel Palade bodies gets secreted towards the apical side into the vessel lumen where it remains tethered and forms ‘sticky’ VWF strings which help to capture platelets, whereas low molecular weight VWF is constitutively secreted basolaterally where it becomes incorporated into the vascular basement membrane (*65*). In contrast, plasma VWF mainly mediates platelet adhesion to exposed subendothelial collagen and gets incorporated into growing thrombi. The contributions of platelet-secreted VWF to these functional categories is not well defined. Platelet-derived VWF is dispensable for haemostasis and arterial thrombosis (*66*) but partially contributes to ischaemic stroke injury (*67*). However, where platelet-derived VWF gets deposited – at the subendothelial matrix, in the thrombus or into the vessel lumen – is difficult to study *in vivo*.

At early stages of platelet spreading, VWF staining localized primarily to the central granulomere region (Figure 7A; ‘o’). The disperse and spatially rather homogeneous VWF staining at the coverslip did not change much across time points and thus probably reflected residual plasma VWF present in the washed platelet preparation. VWF frequently stuck to the lateral sides of the cell body (arrowheads), clearly outside of the cell, indicating it was released and had remained attached to the cell. At later time points, the granulomeric VWF staining remained prominent and VWF was increasingly seen at lamellipodia. Several microns long VWF ‘mini’ strings formed (arrows) which were often attached to granulomeric or lamellipodial VWF pools. 3D STORM showed that these VWF structures were at different heights (Figure 7B). Dual color 3D STORM with GPIbα (CD42b) co-staining further demonstrated that most VWF was localized to the apical membrane or was found beneath the cell (Figure 7C; yellow arrowheads). VWF retention at spreading platelets was >80% on fibrinogen and >90% on collagen over the studied duration of 20 minutes (Figure 7D). We noted that some of the side-ways secreted VWF was not covered by lamellipodia and thus outside of the cell outline (asterisks), which was not considered ‘retained’ by our analysis.

**Figure 7.**
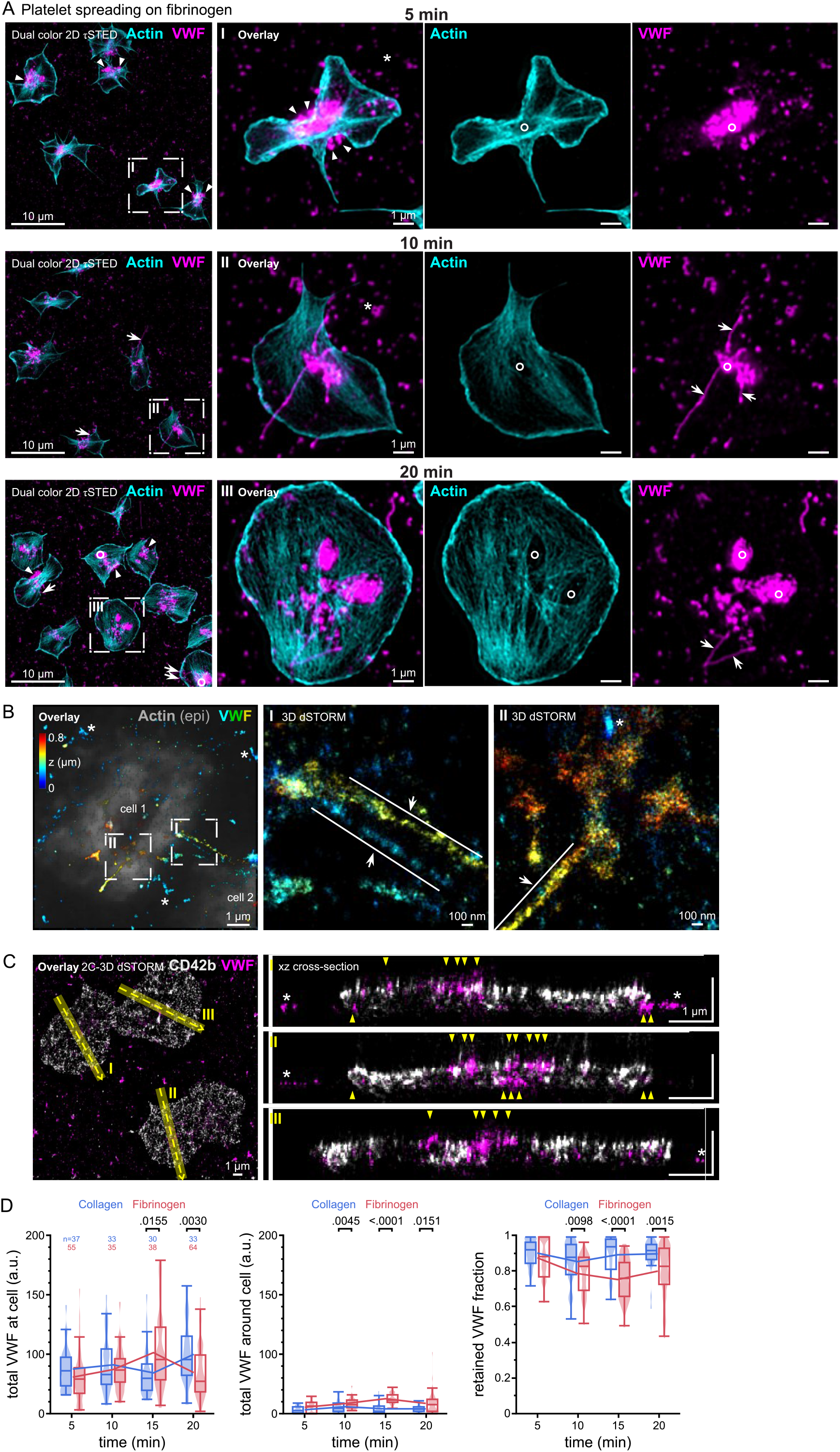
Secreted VWF is retained at apical and basal platelet membranes. Washed human platelets were spread on fibrinogen- or collagen I-coated coverslips up to 60 min, fixed at different time points, permeabilized, immunostained, and imaged by SRM. **A**. Representative overview images and magnified single cell insets of dual-color 2D τSTED f-actin (cyan) and VWF (magenta) images after spreading for 5, 10 or 20 min. Images show regular side-ways secretion of VWF in concave cell regions (arrowheads), originating from the granulomer region which is devoid of f-actin (‘o‘). VWF strings (arrows) are frequently observed. **B**. Single cell 3D dSTORM image of VWF (color-coded according to z-position). The overlay shows f-actin (gray) for context. VWF localizes either close to the coverslip (blue) or on top of the cell (yellow/orange). **C**. Single cell dual color 3D dSTORM image of VWF (magenta) and GPIbα (CD42b; gray) and xz cross-sectional sideviews. The superior z-resolution of dSTORM confirms co-localization of the secreted VWF staining with the plasma membrane (yellow arrowheads), and disperse VWF bound to the coverslip (asterisks). **D**. Total VWF intensity around (left) or at (middle) the cell and their ratio (right). VWF was mostly retained at the cell, moreso for platelets on collagen I compared to fibrinogen.

In summary, spreading platelets secreted VWF into all directions where it remained tethered to the membrane with limited mobility and could be unraveled into VWF strings.

### Strong agonist stimulation overrides cytoskeletal-secretory coordination

Soluble agonists activate resting suspended platelets, inducing the fusion of α-granules with nearby membranes (*10, 11*), as well as contributing to integrin activation and cytoskeletal changes (*18, 30*). In spreading platelets, different agonists also shift the relative activity of Rho family members that govern protrusion formation and platelet contractility (*17, 68*).

Agonist type and concentration matter: for example, high dose thrombin favors the ‘compound fusion’ of α-granules with each other or with the open canicular system in suspended platelets (*10*) and drives RhoA-mediated platelet contractility in adherent platelets (*68*). We thus asked whether different agonists perturb the coordination between secretion and cytoskeletal reorganization in spreading platelets.

In the presence of agonists, platelets on fibrinogen secreted and spread faster, having completed these processes after 20 minutes (Figure 8A, Supplementary Figure S8 A+F). Noticeable differences were seen between agonists at early time points in terms of P-selectin expression and filopodia formation. Platelets exposed to high-dose thrombin (1 U/mL; THR-1) had fully secreted already after 2 minutes, whereas platelets spreading in the presence of PAR4-activating peptide (PAR4AP; 100 µM) or CRP (5 µg/mL) required 5 or 10 minutes to degranulate, respectively (Figure 8B). Significant transient filopodia formation was induced by CRP at early time points, and to a lesser extent by PAR4AP, whereas it was strikingly suppressed by high-dose thrombin (Figure 8C). Lamellipodia formation and spreading dynamics were comparable between agonists (Figure 8D). The effects of the PAR1 agonist TRAP6 (10 µM) were comparable to CRP and PAR4AP, whereas ADP (5 µM) or low-dose thrombin (0.1 U/mL; THR-0.1) induced slower degranulation and lesser filopodia (Supplementary Figure S8 B-E).

**Figure 8.**
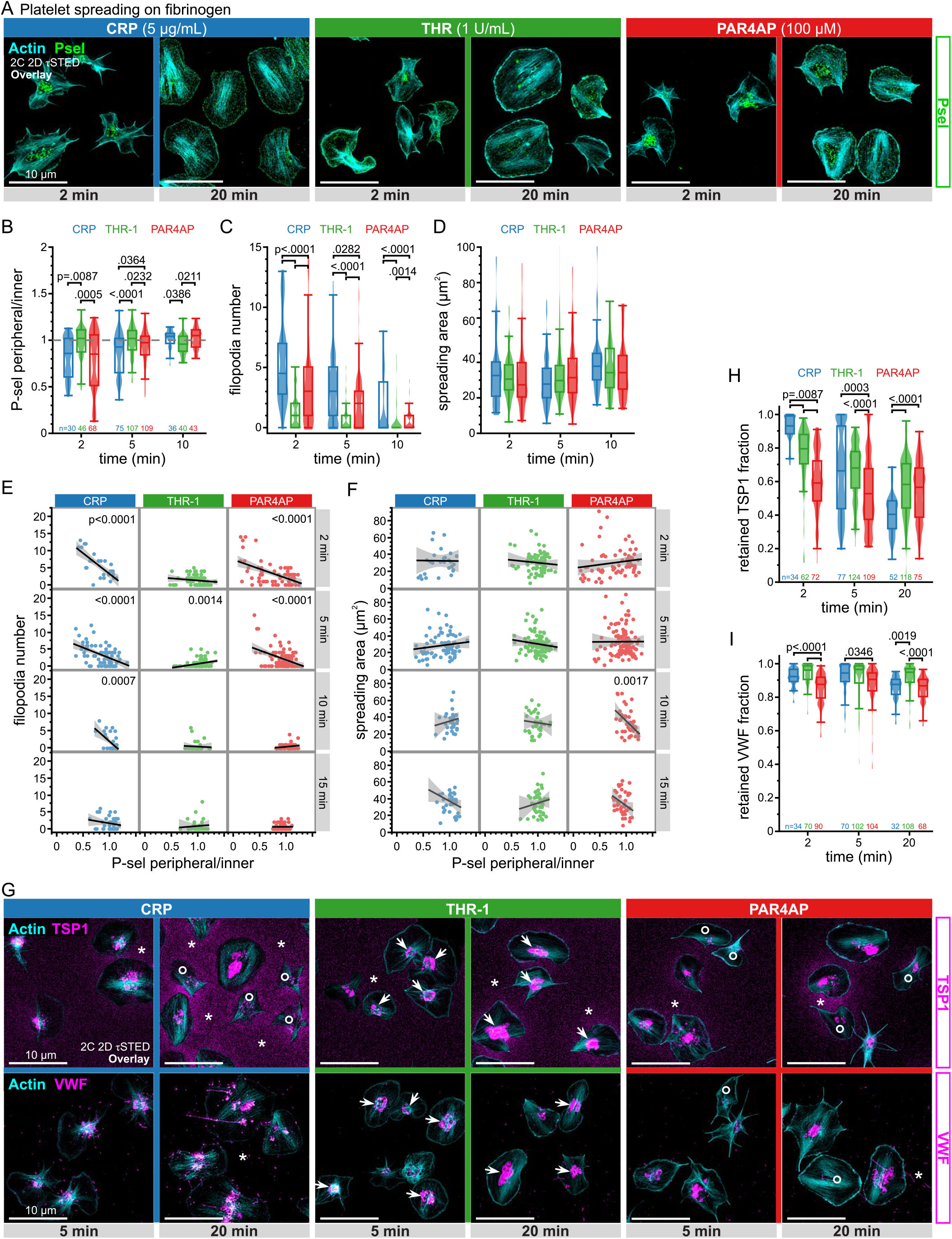
Strong agonist stimulation overrides cytoskeletal-secretory coordination. Washed human platelets were spread in the presence of agonists on fibrinogen-coated coverslips up to 20 min, fixed at different time points, permeabilized, immunostained, and imaged by2D τSTED. **A**. Representative SRM images of f-actin (cyan) and P-selectin (green) in human platelets spread on fibrinogen for 2 or 20 minutes in the presence of 5 µg/mL CRP (blue; left), 1 U/mL thrombin (green; middle) or 100 µM PAR4-activating peptide (PAR4AP; red; right). Scale bars: 10 µm. **B**-**D**. Morphometrics of platelets spread in the presence of different agonists (legend) for 2-10 minutes. Single-cell parameters are: **B**. Peripheral-to-inner P-selectin intensity ratio; **C**. Filopodia number; **D**. spreading area. Single cell data are shown as violin plots and summarised as box plots (25/50/75 percentile, whiskers excluding outliers). Indicated in (B) is the total number of platelets per time point pooled from 1-3 independent experiments. Statistical tests by ANOVA with Tukey for multiple comparisons. THR-1: thrombin at 1 U/mL. **E**-**F**. Correlation analyses of **E**. filopodia number or **F**. spreading area versus the degree of α-granule secretion (measured by the peripheral-to-inner P-selectin intensity ratio) at early time points. An anti-correlation of the filopodia number is observed for CRP and PAR4AP but not for THR-1. No significant correlations were seen for the spreading area in the presence of agonists. **G**. Representative τSTED images of f-actin (cyan) and TSP1 (magenta, top row) or VWF (magenta, bottom row) in human platelets spread on fibrinogen for 5 or 20 minutes in the presence of agonists as in (A). Scale bars: 10 µm. **H**-**I**. Fraction of **H**. TSP1 or **I**. VWF retained at the cell compared to its immediate surrounding (for definition, see Figure 6C) for platelets spread in the presence of different agonists. Visualisation and statistical tests as in (B). The total number of platelets per time point pooled from 1-3 independent experiments is denoted below the graphs.

While platelets underwent agonist-supported degranulation, their filopodia number decreased with progressing P-selectin membrane expression, whereas this correlation was lost when secretion reached completion (Figure 8E). This was true for all agonists except ADP where the correlation did not reach significance (Supplementary Figure S8H). The degree of spreading in the presence of agonists showed no systematic correlation to α-granule secretion (Figure 8F, Supplementary Figure S8I), in contrast to spreading in the absence of soluble agonists (see Figure 5H). In summary, agonists accelerated secretion dynamics and abolished the dependence of spreading on degranulation, while concurrent filopodia formation remained temporally correlated with degranulation.

The altered degranulation dynamics also affected the fate of α-granule contents. Secreted TSP1 accumulated prominently around CRP-stimulated platelets (asterisks), less so for high-dose thrombin or PAR4AP-stimulated platelets (Figure 8G). While some CRP- or PAR4AP-stimulated platelets were nearly devoid of TSP1 (‘o’), high-dose thrombin led to a pronounced central fused appearance of TSP1 (arrows). As a result, retention of TSP1 was highest in thrombin- and lowest in CRP-stimulated platelets, with PAR4 stimulation in-between (Figure 8H). TRAP6 had comparable effects to PAR4AP, while ADP and low-dose thrombin showed high retained TSP1 fractions (Supplementary Figure S8 J+K) in agreement with their slower secretion dynamics. Secreted VWF showed a qualitatively similar behavior as for TSP1, but at much higher overall cellular retention when comparing CRP, high-dose thrombin and PAR4AP (Figures 8 G+I) or other agonists (Supplementary Figure S8 J+L). Of particular note, high-dose thrombin, and to some extent also low-dose thrombin, led to a prominent centralization of VWF and fewer VWF strings at lamellipodia (Figures 8 G, Supplementary Figure S8 J). Taken together, platelets showed signs of compound fusion in response to high thrombin concentrations, limiting the escape of granule contents or their display at the platelet surface.

## DISCUSSION

In this study, we combined τSTED microscopy, 3D dSTORM, quantitative image analysis and computational simulations to investigate the spatial organization and secretion dynamics of platelet α-granules. We show that major α-granule constituents occupy distinct nanoscale subdomains, with P-selectin, TSP1 and VWF exhibiting characteristic and largely non-overlapping morphologies within intact granules (Figures 1-3). During platelet spreading, intact α-granules remained confined to the central granulomere and were absent from filopodia and lamellipodia, identifying the platelet cell body as the exclusive site of α-granule exocytosis (Figure 4). Quantitative single-cell analyses revealed that degranulation is temporally associated with the transition from filopodia-rich to lamellipodia-rich morphologies (Figure 5), although experiments using different adhesion substrates and soluble agonists indicate that these processes are coordinated by overlapping signaling pathways rather than direct spatiotemporal coupling (Figures 5+8). Finally, we demonstrate that α-granule cargoes follow distinct extracellular trajectories after secretion: TSP1 accumulates within diffusion-restricted pockets beneath platelets and in the surrounding matrix, whereas VWF remains highly retained at platelet membranes and can form membrane-associated strings (Figures 6-8). Together, these findings provide a revised framework for understanding how platelet α-granule organization, secretion and cargo retention shape the local platelet microenvironment (Figure 9).

**Figure 9.**
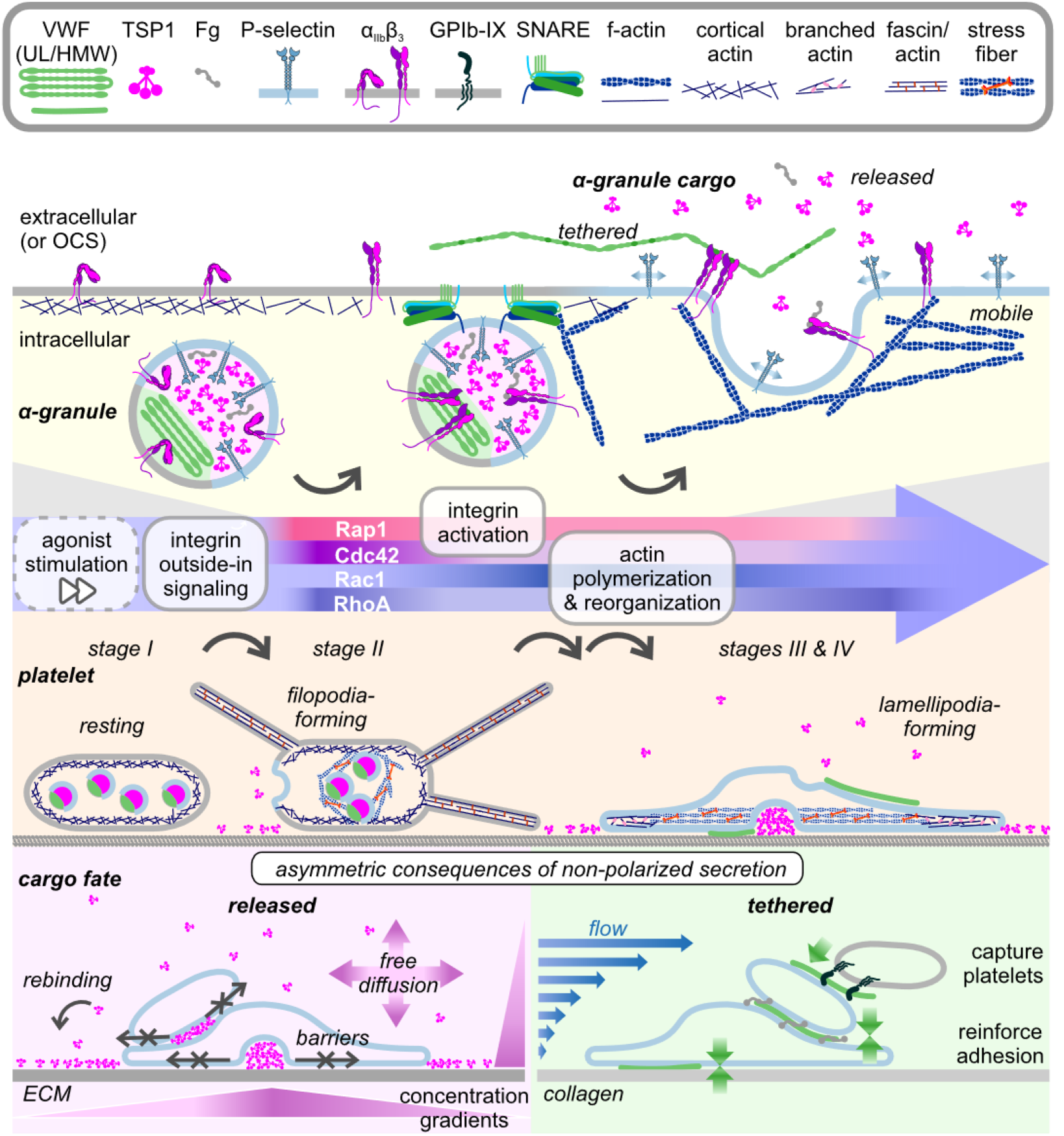
Platelet biomechanics and secretion shape the fate of α-granule cargo. Schematic summary of results. Upper panel: VWF and TSP1 are contained in speparate sub-compartments of α-granules, with P-selectin being excluded from VWF-proximal membranes. Granule exocytosis results in mobile P-selectin, tethered VWF forming strings, and released TSP1. Middle panel: Platelet spreading proceeds through subsequent filopodia and lamellipodia formation driven by major actin rearrangements (middle). The central α-granule pool gets secreted mainly between stage II to III, without directional bias. Agonists shift the Rho signaling balance, differentially accelerating spreading and secretion. Bottom panel: Even though α-granule secretion is not biased, apical and basal secretion serve distinct purposes. Left: Released TSP1 might diffuse and bind to extracellular matrix (ECM) or encounter barriers and accumulate beneath platelets, effectively creating 3D concentration gradients. Right: Tethered unwound VWF might reinforce platelet-collagen adhesions on the basal side, mediate apical platelet recruitment, and reinforce platelet-platelet interactions.

Our data clearly show that granule contents, here TSP1 and VWF, are not distributed across the entire granule internal volume but rather occupy sub-compartments. This was known for VWF (*41, 51*) but is not usually considered for other components. The concept of co-localization, i.e. whether two components reside in the same space, is thus not appropriate to judge if these components co-occupy, or are contained within, the same granule. The better the optical resolution is relative to the size of α-granules, the more pronounced this effect becomes, but it is expected to bias analyses already at confocal resolution. The close packing of granules in resting platelets (*50*) poses additional challenges, necessitating a delineation in 3D of what is an individual granule. A rigorous SRM approach to study the compositional heterogeneity of α-granules thus would require SRM 3D imaging data of a reliable α-granule membrane marker that can be used for segmentation of individual α-granules, and the simultaneous measurement of two (or more) components. Our finding that the P-selectin staining was also not evenly distributed around the surface of α-granules limits the ability to distinguish individual granules based on this staining. Our computer simulations showed that the uneven P-selectin staining of experimental images is not sufficiently explained by random factors alone. Additional reasons could be of technical or biological nature. Hindered access to the P-selectin antibody epitope, which resides on the inside of the densely packed α-granule, could be one reason. Exclusion of P-selectin from certain granular membrane regions due to the clustering of other membrane proteins could be another reason. Our discovery that the P-selectin staining was suppressed close to VWF is compatible with either explanation. VWF is stored densely packed in peripheral compartments of α-granules close to their membrane (*41, 51, 69*), similar to its tubular storage in Weibel Palade bodies (*70*). On the other hand, the α-granule membrane contains adhesion proteins such as integrins and GPIb, which can bind to VWF. We thus speculate that these could be clustered in vicinity of stored VWF, hindering access of P-selectin to these membrane compartments. Upon α-granule fusion with the plasma membrane, these adhesion proteins could hypothetically also mediate the retention and anchorage of VWF at the platelet surface, as observed in our experiments.

Thanks to its superior resolution compared to diffraction-limited microscopy, SRM reliably distinguished between P-selectin staining of intact granules and disperse P-selectin membrane expression which had distinct morphologies. This enabled us to pinpoint the sites of α-granule exocytosis in spreading platelets to the central part of the cell body containing the granulomere. To transport α-granules through thin and actin-filled lamellipodia or filopodia, these structures would need to widen to accommodate the thicker α-granules. The most likely explanation for a peripheral localization of P-selectin and other proteins that mediate degranulation (including VAMP7) (*13, 29*) thus is that these proteins diffused or were transported there after fusion of granules with the plasma membrane or when not bound to granules. For example, it is plausible that α-granules fuse with the plasma membrane when this side of the cell body flattens during lamellipodia formation, and that granule-associated proteins remain membrane-bound and become integrated into the lamellipodium and thus move outwards (*29*). Alternative explanations for apparently divergent findings could be that platelet spreading on the non-physiological ligand poly-L-lysine (*13, 29*) proceeds differently than integrin signaling-mediated spreading on natural adhesion proteins, e.g. forming thicker lamellipodia, or that endocytosed cargo(*29*) might be stored in α-granules of smaller size that can more readily be transported to the periphery.

Our data do not indicate a preference or bias for α-granule secretion into a certain direction. This conclusion is supported by several observations. Firstly, disperse P-selectin staining was localized to basolateral and apical membranes to similar extents. Although diffusion along the continuous plasma membrane may average out potential differences over extended times, no biased localization was observed at shorter times. Secondly, secreted TSP1 accumulated both beneath the cell as well as around the cell. Although the relative staining intensity of TSP1 within/beneath versus around spread platelets is only a semi-quantitative measurement which does not strictly separate basal versus apical secretion, the values of ∼0.3-0.6 on average after extended times indicate that there was no systematic directional preference (=0.5). Thirdly, secreted VWF remained tethered to basolateral and apical platelet membranes and was also observed at lateral sides of the cell body during early phases of spreading. An exclusive basolateral secretion from adherent platelets (*14*) would require a defined cellular polarity. In epithelial cells, the exocyst complex delivers proteins to the basolateral membrane, thereby establishing polarity of secretion (*71*). The exocyst complex also supports dense granule secretion in activated platelets under the control of the membrane tethering Ras-like GTPase Ral (*72*). However, it is questionable if this mechanism could establish a polarity within seconds to minutes, which is the relevant time scale for granule secretion after platelet adhesion, whereas it requires 3-6 hours in epithelial cells (*73*). Studying Ral sub-cellular localization could be an opportunity to obtain further insights into a potential polarity of adhered platelets, a largely unexplored topic. If platelets had the ability to direct their secretion into different directions, this could have profound consequences for understanding the establishment of thrombus architecture, but experimental evidence is lacking.

Even without showing signs of polarized secretion, our data confirm and extend the concept of self-deposition of adhesion proteins that allows platelets to bridge non-adhesive environments (*14*) or build up a thrombus. Secreted proteins that do not re-bind will diffuse and stick to nearby areas of the environment, as observed for fibrinogen (*14*), or here for TSP1. The laws of diffusion dictate that the highest concentration will be close to the source, leading to the deposition in form of a rim around platelets. In case of platelets spreading on bare glass (*31*) or on adhesive islands (*14*), secreted adhesion proteins such as fibrinogen thus will enable further platelet spreading, migration, or adhesion of other platelets over areas that initially were non-adhesive. The decreasing surface concentration of these granular proteins away from the secreting platelet naturally limits the effect to their immediate vicinity. In a similar manner, secreted growth factors are expected to form short-range gradients throughout a thrombus or at wound sites, providing cues for the invasion by immune cells (*74*), in addition to adhesive tracks in the immediate vicinity of platelets, which include platelet-produced fibronectin fibrils (*33*).

In an extension to this concept, we here observed that VWF is retained at platelet membranes. The extent of VWF retention was very high (>80%) and consistent with findings from agonist-stimulated washed platelets in suspension (*52, 75*). It was consistently high even in the absence of any apparent compound granule fusion, and VWF localized to the membrane rather than intracellularly. A very efficient surface retention, which has been shown to be dependent on integrin αIIbβ3 (*75, 76*) either requires very effective rebinding, i.e. very sticky VWF molecules, and/or an active tethering during secretion. Integrins in the granule membrane (*77*) face the granule interior and transition to a binding-competent state following initial platelet adhesion, as expected from integrin inside-out signaling because their tails face the cytosol. We thus speculate that integrin adhesion receptors might already bind adhesion proteins stored within α-granules, such as VWF, and retain this connection during granule fusion with the plasma membrane, effectively tethering secreted adhesion molecules at the platelet surface. This proposed mechanism could support the related hypothesis that platelet VWF is prebound to receptors after granule fusion to the open canicular system(*76*).

In our experiments, tethered VWF partially became elongated. This extended conformation of VWF is known to support platelet-VWF binding (*78*) and could mediate the recruitment of additional platelets to form a thrombus(*76*). The capture of platelets at the exposed subendothelium or at the luminal side of a growing thrombus is usually mediated by plasma VWF, as apparent from normal hemostasis in the absence of platelet VWF (*66, 79*). However, platelet VWF can at least partially compensate for the lack of plasma VWF (*80*), indicating that platelet VWF contributes to, but is not strictly required, for hemostasis (*77*). Basal secretion and retention of platelet VWF might potentially act as a glue to reinforce the anchorage of platelets to collagen(*81*), whereas apical tethering could help to recruit platelets to a growing thrombus, similar to VWF strings at the apical surface of activated endothelial cells. In the context of thrombosis, pre-tethered and unfolded VWF at the surface of activated platelets could potentially initiate the formation of VWF-platelet aggregates. This possibility is particularly intriguing because platelet-derived VWF has been shown to aggravate tissue injury post-ischemic stroke (*66*), and VWF-platelet aggregates are critical to the ‘no-reflow’ phenomenon, preventing successful reperfusion after mechanical thrombectomy (*82*). The formation of these aggregates requires perturbed flow conditions and is sensitive to ADAMTS13, suggesting the involvement of hemostatically active ultra-large (UL) or very high molecular weight (HMW) VWF multimers (*82*). Platelet-derived VWF is enriched in UL-VWF compared to circulating VWF(*51, 83*) because α-granule storage protects it from proteolytic degradation (*84*). Moreover, several stimuli can activate platelets during ischemic stroke, including perturbed flow dynamics and endothelial activation (*85*). Taken together, it seems plausible that platelet-tethered VWF after platelet activation could contribute to the no-reflow phenomenon in infarcted areas.

Our identification of TSP1-containing ‘pockets’ beneath spread platelets highlights that the fate of released α-granule cargos depends strongly on whether they encounter local barriers. While the local accumulation of TSP1 observed in our experiments resulted from the close apposition of platelets to the coverslip, similar diffusion-restricted microenvironments may occur when platelets become incorporated into densely packed thrombi or seal vascular microlesions. In these situations, released cargo may attain markedly different concentrations on the basal and apical sides, leading to substantially greater effects on neighboring cells exposed to the basal compartment. This conceptual interpretation is supported by the diffusion profiles across the core-shell thrombus model(*86*) that help to limit thrombus growth.

Platelets spreading on fibrinogen or collagen in the absence of soluble agonists showed the highest secretory and cytoskeletal activity over the first 10 minutes. The correlations detected on the single cell level are consistent with filopodia formation preceding degranulation preceding lamellipodia formation, and platelets within the population undergoing this sequence at different speeds. This sequence is in agreement with live-cell observations (*87*) and the accepted model for morphological spreading stages I-IV (*88*), as well as with stochastic secretion kinetics (*12*). The question is whether these processes are coordinated separately, they share regulatory mechanisms, or even are dependent on each other.

It is generally accepted that α-granule secretion enables spreading by providing additional membrane for protrusion formation (*13, 29*), beyond additional membrane from the open canicular system. The fact that degranulation precedes lamellipodia formation is compatible with this hypothesis but does not prove causation. The observation that the area after 60 minutes of spreading on fibrinogen, when spreading is completed but not all cells have secreted their granules, did not correlate positively with P-selectin membrane expression means that incomplete secretion does not restrict spreading. Platelet spreading on uncoated glass (*89*) or over non-adhesive areas (*14*) requires self-deposition of adhesion proteins and thus depends on α-granule secretion, which makes it hard to distinguish from the provision of α-granular membrane. Our observation that lateral secretion onto an already adhesive substrate does not necessarily lead to lamellipodia (or filopodia) formation at the secretion site speaks against a strict local intimacy. This is true not only for the membrane provided by an α-granule, but also for a hypothetical switch of VAMP7 from mediating granule fusion to increasing Arp2/3 activity locally at secretion sites (*13*). While the importance of VAMP7 is well-documented for either process (*13, 29*), our data do not indicate that these processes are limiting each other through the shared VAMP7 pool.

Soluble platelet agonists quite universally accelerated granule secretion in platelets spreading on fibrinogen, with a similar hierarchy as in suspended platelets (*12*) and also accelerated spreading dynamics. These outcomes are well understood to be the consequence of increased G-protein coupled receptor signaling converging on Rap1 activation which on one hand induces α-granule secretion and on the other hand enhances integrin inside-out activation and, indirectly, subsequent outside-in signaling responsible for driving lamellipodia formation (*30*). Shared signaling paths are thus a sufficient and plausible explanation for the observed correlation. When comparing platelets spreading on collagen versus fibrinogen, a higher filopodia number on collagen correlated with a higher degree of degranulation at the early 2-minute time point, whereas this was not the case in the presence of soluble agonists which had differential effects on degranulation and filopodia formation.

These findings exemplify the differences in integrin-initiated versus agonist-induced pathways leading to α-granule secretion (*30*) or filopodia formation, with different adhesion proteins and soluble agonists shifting the balance between these pathways. Again, regulatory pathways overlap, namely on Cdc42 (*15, 90, 91*), which could explain correlations without the necessity for a direct interdependence.

Interestingly, major actin stress fibers were never crossing the location of intact α-granules nor the sites of their basal secretion, which are identifiable by the ’pockets’ of retained TSP1 beneath the cell. One possible explanation is steric hindrance, i.e. that the space was pre-occupied by granules or the membrane pocket. Another contributing factor could be that surface receptors in this pocket are bound to secreted adhesion proteins which remain mobile, thus preventing the mechanosensory feedback that is necessary for focal adhesions formation and maturation to develop in this area. Whether other mechanisms on the signaling level, for example a spatial separation and antagonistic activity of RhoA and Rac1 (*92*), contribute to the development of this mutual exclusion remains subject to speculation.

Taken together, our data do not support a spatiotemporal intimacy and interdependence between α-granule secretion on one side and actin cytoskeletal rearrangements on the other side, but are sufficiently explained by overlapping regulatory pathways and a mutual spatial exclusion between α-granules or their secretion pockets and major f-actin stress fibers.

The improved resolution and complementary usage of two SRM techniques, complemented by computer simulations to rationalize image interpretation and single cell morphometric analyses to quantify cellular behaviors are key strengths of our study, but our approach also had several limitations. (i) Technical limitations. Temporal correlations were derived from population data of fixed cells; live-cell SRM (*93*) could provide further support for our interpretation, but typically at a compromised optical resolution and restricted choice of staining targets. Fixation prevented the selective staining of membrane-expressed P-selectin because fixation compromises membrane integrity, whereas efficient f-actin staining required membrane permeabilization. The non-observation of TSP1 at platelet membranes was surprising given its well documented binding affinity for CD36 (*94*); potential explanations could be that CD36 binding masks the antibody epitope on TSP1, or that the permeabilization procedure partially solubilized CD36 even after fixation. We thus cannot exclude that a portion of TSP1 was bound to the membrane but not detected. Fast diffusion and equilibration of membrane-expressed P-selectin could have hampered the identification of granule fusion and protrusion events happening in quick succession. The peripheral-to-inner P-selectin ratio is overestimated in less spread platelets due to an imperfect boundary and the higher height of platelets, which is not fully captured by a single 2D τSTED image. The ratios used to measure TSP1 and VWF retention have the advantage of reducing the dependence on unreliable absolute fluorescence intensity measurements, but do not account for granule cargo being washed away, thus potentially overestimating the proportions retained within/beneath the platelet; the derived ratios are valuable for relative comparisons but their absolute values shouldn’t be overinterpreted. (ii) Experimental limitations. Using washed platelets, spreading on continuous adhesion protein coatings, and selected soluble agonists are well established models to study single platelet behavior, however, these choices strategically simplify the much more complex context of platelet adhesion at injury sites in terms of dimensionality, protein composition, fluid shear, platelet-platelet interactions, different agonist concentrations and combinations. Our findings thus were interpreted in this *in vitro* context, and their projection towards pathophysiological scenarios *in vivo* needs to account for the differences. Due to the overlap in signaling pathways controlling degranulation and cytoskeletal changes, a selective targeting of e.g. α-granule secretion alone without affecting other cytoskeletal processes was not possible but would be of high interest to obtain additional insights. Despite these limitations, our approach resolved granular organization with unprecedented detail, clarified questions on secretion polarity, and delineated the extent to which α-granule secretion and cytoskeletal dynamics could influence each other.

## MATERIALS AND METHODS

### Materials

Adenosine diphosphate disodium salt (#A2754), apyrase from potato (#A6535), bovine serum albumin (#A7906), CF 660C succinimidyl ester (#SCJ4600050), CF 680 succinimidyl ester (SCJ4600055), catalase, Cysteamine (M9768), collagen type I from calf skin (#C9791), 1,4-Diazabicyclo[2.2.2]octane(DABCO, D27802), D-glucose (#G7528), fibrinogen from human plasma (#F3879), glucose oxidase(#G2133), Mowiol® 4-88 (#81381), prostaglandin E1 (#P5515), potassium chloride, potassium phosphate, paraformaldehyde (PFA), sodium chloride, sodium citrate, sodium phosphate dibasic, thrombin (T6884), Triton™ X-100 were purchased from Merck. Mouse anti P-selectin (clone AK4; # SC19996 L) was from Santa Cruz Biotechnology. Abberior^®^ STAR 635P NHS ester (#07679), STAR 635 phalloidin (#30972), Alexa Fluor (AF) 488 NHS ester (#A20000), AF 488 phalloidin (#A12379), AF 594 NHS Ester (#A20004), AF647 NHS ester (#A20006) AF 647 phalloidin (#A22287), mouse monoclonal antibodies against thrombospondin-1 (TSP-1; Clone A6.1, #MA5-13398) and CD42b (#MA5-16565) were from Invitrogen. Unconjugated donkey anti mouse IgG (#715-005-150), donkey anti rabbit IgG (#715-005-152) and donkey anti goat IgG (#705-005-147) antibodies were from Jackson ImmunoResearch. TRAP-6 (PAR1-AP, SFLLRN; #HY-P0078) and PAR4-AP (AYPGKF; #HY-P1309) were from MedChemExpress. Mouse anti thrombospondin-1 (TSP-1; Clone A6.1) and rabbit polyclonal antibody against CD41 (ab134131) were from Abcam. Ethylenediaminetetraacetic acid (EDTA) tube (S-Monovette, 3mL) and citrate blood collection tubes (S-Monovette, 10 ml) were from Sarstedt. Others: CRP-XL (CambCol Laboratories); coverslips (20 mm diameter, thickness 1.5; Hecht-Assistent, Germany); eptifibatide (Integrilin©; GlaxoSmithKline); Goat anti TSP-1 antibody (R&D systems, #AF-3074); Rabbit polyclonal anti-VWF antibody (Dako; Agilent); 23 nm Abberior Star635P GATTA-Bead R (GattaQuant GmbH).

### Blood collection and platelet preparation

Ethical approval was sought from the RCSI Research Ethics Committee (#1394 and #1504) before commencement of the study. All protocols agreed with local legislation and the Declaration of Helsinki. Written informed consent was sought from healthy volunteers. Phlebotomy was performed using citrate blood collection tubes. Washed human platelets were prepared following established protocols (*95*).

### Resting platelet sample preparation

Cleaned 20 mm diameter round glass coverslips (precision thickness #1.5) were coated with fibrinogen (0.1 mg/mL) in PBS. Washed platelets (10 × 10^6^/well) were fixed with 3% paraformaldehyde (PFA) for 10 minutes, transferred onto fibrinogen-coated coverslips, centrifuged at 1000 RCF for 10 min (acceleration 5, deceleration 5), and then quenched with Tris buffer.

### Platelet spreading on fibrinogen/collagen

Cleaned 20 mm glass coverslips were coated with fibrinogen (0.1 mg/mL) in PBS or collagen type I in 0.1% acetic acid for 2 h at room temperature and washed three times with PBS. Washed platelets (10 × 10^6^/well) were added to coated coverslips in the presence or absence of agonists (5 µM ADP, 5 µg/mL CRP-XL, 10 µM TRAP-6, 100 µM PAR-4, or 1 U/mL thrombin) and incubated at 37°C and 5% CO₂ in Tyrode’s buffer. After 2 min, non-adherent platelets were removed by aspiration. For longer time points, fresh Tyrode’s buffer containing the respective agonist was added, and incubation was continued for a total of 5, 10, 20, or 60 min. Samples were then washed with Tyrode’s buffer and fixed with 3% PFA in PBS for 15 min at room temperature.

### Fluorescence stainings

Samples were washed with PBS, permeabilized with 0.5% Triton X-100 for 15 min, and blocked with 3% BSA in PBS for 1 h at room temperature. For τSTED, platelets were stained for the α-granule markers P-selectin (1:100), VWF(1:200), and TSP-1 (1:100) using primary antibodies overnight at 4°C, followed by the corresponding secondary antibodies conjugated to Alexa Fluor (AF) 594 or Abberior® STAR 635P (degree of labelling 3–5) for 4 h at room temperature. F-actin was stained with phalloidin-AF 488 (1:100; for confocal) or Abberior STAR 635P phalloidin (1:40; for τSTED), in parallel to secondary antibody incubation. For dSTORM, fixed spreading platelets were stained for the α-granule markers P-selectin, TSP-1, and VWF, and the surface receptors GPIbα (1:100) or integrin α_IIb_ (1:100) using primary antibodies followed by AF647- or CF680-conjugated secondary antibodies (degree of labelling 1–2). Following final washes, samples for τSTED were mounted on glass slides using Mowiol® 4-88 containing 4% DABCO, whereas samples for dSTORM were kept at 4°C in PBS containing 0.05% sodium azide.

### τSTED microscopy

τSTED imaging was performed using a Leica Stellaris STED microscope (Leica Microsystems, Germany) with 594 nm and 638 nm white-light laser excitation (25% and 15% intensity, respectively, for resting platelets; 15% intensity each for spread platelets), a 775 nm depletion laser (30% intensity), and a 100× oil immersion objective (HC PL APO CS2, 1.40 NA). Images of resting platelets were acquired at a pixel size of 0.016 µm or less (4× zoom or higher). Images of spread platelets were acquired at a pixel size of 0.021 µm (3× zoom). Lifetime was gated 0.5-8 ns with tau-contrast set to 120 strength and 60 denoise.

### Morphometric analysis of spread platelets

Morphometric analysis of the platelet actin cytoskeleton and α-granule secretion was performed in Matlab (R2024a; MathWorks) by extending a previously validated approach (*95*). Briefly, f-actin images were used to segment individual platelets. From the cell outline, parameters including platelet spreading area, circularity and filopodia number were extracted. The automatic detection of filopodia was implemented by smoothing the outline coordinates with a fine and a coarse moving window. The coarser outline was subtracted from the finer smoothed outline to determine maxima, minima and intersection points. The latter were moved by a predefined fraction towards the adjacent minima. The corresponding coordinates of the raw outline for maxima (=tip of protrusion) and intersection points (=base of protrusion) were determined. The distance between the middle of the base intersecting line and the tip was taken as the length of the protrusion. The width of the protrusion was taken as the vector component of the intersecting line perpendicular to the protrusion direction. Protrusions with an aspect ratio length/width ≥1.2 and lengths ≥0.45 µm were counted as filopodia. From the f-actin stain, the internal alignment of actin filaments was determined. To determine the P-selectin intensity ratio between peripheral to central regions, a distance map was created from the cell outline, and taking the distance yielding sub-areas of same size, cumulative staining intensities of the peripheral *I_peri_* and central *I_inner_* regions were used to calculate the intensity ratio as 2 × *I_peri_*⁄(*I_peri_* + *I_inner_*). For determining the retained fraction of TSP1 or VWF, the cell outline was dilated by 1 µm, excluding intersections with nearby cells, and the cumulative staining intensities within the cell area *I_cell_* and within the surrounding rim *I_rim_* were used to calculate the intensity ratio *I_cell_*⁄(*I_cell_* + *I_rim_* ).

### dSTORM microscopy

Sample coverslips were mounted in a magnetic imaging chamber and immersed in super-resolution imaging buffer containing 1 mg/mL glucose oxidase type VII, 0.2 mg/mL catalase, 90 mM cysteamine, and 4% (w/v) glucose. Single molecule imaging was performed on a custom-build microscope(*96*). Regions of interest were selected and focused. Up to 120,000 frames (if blinking did not cease earlier) in two simultaneous channels were acquired at 50 s^-1^ using 647 nm laser excitation, a longpass beamsplitter (665 nm or 690 nm), and two 700/100 bandpass emission filters. Three-dimensional single-molecule localisation was enabled using an astigmatic lens common to both channels. A 405 nm laser with gradually increasing intensity was applied to promote fluorophore recovery from the dark state and was activated automatically only when the number of blinking events dropped below a predefined threshold.

### dSTORM visualization and analysis

All data processing for 3D dSTORM data was done in SMAP (*97*). Spots in images were fitted using a spline PSF. Localizations were assigned to AF647 and CF680 based on ratiometric intensity measurements after determining a spatial transformation between the measured channels (split by wavelength). z-coordinates were corrected for the refractive index mismatch between immersion medium (oil) and sample medium (buffer) by multiplication with a constant factor of 0.75. Chromatic offsets between channels were corrected by adding a constant offset to CF680 which was measured using a dual-labeled coverslip coating. Localizations were grouped and filtered according to loglikelihood (>-0.8) and localization precision (<20 nm). Visualization used Gaussian rendering with the localization precision as width and a 10 nm pixel size. Measurements of the filopodia width were done manually in Fiji by drawing lines on rendered images (dSTORM) or masks of cell outlines (τSTED). For automated measurements of the lamellipodia height from 3D dSTORM data of CD41 stainings, sites along the cell edge were picked in SMAP’s Site Explorer. A custom written evaluation plugin oriented the sites along a line that was drawn perpendicular to the cell edge, retrieved all localizations 1 µm to both sides of the line, binned their z-positions and determined the distance between the two main peaks from a smoothing spline, yielding the average lamellipodia height. The plugin also rendered a xz cross-section using 20 nm pixel size.

### Simulated STED images of α-granular protein stains

α-granules were simulated as spheres with diameter *d*, total number *n* and size distribution *P*(*d*) per platelet. To represent the typical 50-80 granules per platelet spanning 200-500 nm in diameter (*22*), we chose *n* = 65 and modelled the size distribution by a normal distribution *P*(*d*) = *N*(µ, *σ*) with µ = 350 *nm* and *σ* = 75 *nm*. The total number of proteins per platelet *N_tot_* was distributed among granules of different sizes as follows: we assumed that membrane markers (P-selectin) were distributed proportional to the granules’ surface areas *S*(*d*), and that inner cargo (TSP-1) was distributed proportional to the granules’ volumes *V*(*d*). The expected number of proteins per granule *Ñ* thus is given by *Ñ*(*d*) = *f*(*d*) · *N_tot_*⁄*n* with the rescaling factor *f* given by *f*(*d*) = *S*(*d*)⁄〈*S*〉 or *f*(*d*) = *V*(*d*)⁄〈*V*〉 for surface markers or inner cargo, respectively, where brackets denote the average of surface (volume) over the granule size distribution 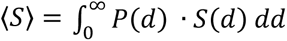. To account for biological stochasticity, the actual number of proteins per granule *N* was randomly drawn from a binomial distribution *N* ∈ *B*(*N_tot_*, *p*(*d*)) with the probability *p* given by this granule’s surface area (or volume) relative to the total surface area (volume) of all granules in the cell *p*(*d*) = *S*(*d*)⁄(*n* · 〈*S*〉). The non-perfect labelling efficiency *ε* of immunostainings was modelled by drawing the actual number of labels per granule *N_L_* from a binomial distribution *N_L_* ∈ *B*(*N*, *ε*). The labels were distributed randomly over the sphere surface (P-selectin, r=d/2) or within the sphere volume (TSP-1, r=0…d/2). Linkage errors were not considered. Each label was conjugated to an average number (degree of labelling, DOL) of 5 fluorophores; to account for conjugation variability, the actual number *N_F_* was randomly picked from the binomial distribution *N_F_* ∈ *B*(10,0.5). Each fluorophore had an expected brightness of ξ̃ = 5 photons per pixel when completely in focus/centre. The expected number of photons sampled at 21.35 nm distances in xy (=pixel size) at a certain focal plane z was calculated by convolving the 3D distribution of the fluorophores with the effective 3D PSF of the τSTED system. The PSF was measured using 40 nm fluorescent beads by acquiring a z-stack with the same acquisition settings as for resting platelet imaging, and then interpolated by a 3D spline (*98*). Alternatively, the PSF was approximated by a 3D Gaussian with lateral width *σ_x_* = *σ_y_* = 35 *nm* and axial extension *σ_z_* = 350 *nm*, which yielded very similar results (not shown). Slices containing the expected photons counts were lastly subjected to Poisson noise to yield the final simulated images.

### Statistical analysis

Plots/statistical analyses of data were prepared/performed in JMP. Plots show mean ± s.e.m. Statistical testing used one-way ANOVA with Tukey’s correction for multiple comparisons. Only comparisons with significance levels <0.05 are shown. Additional data visualization and fitting was performed in Matlab (R2024a; MathWorks).

## ACKNOWLEDGEMENTS

STED images were acquired in the RCSI Super Resolution Imaging Consortium (SRIC) funded by Science Foundation Ireland (18/RI/5723).

## Author contributions

Conceptualization: I.S.; Methodology: S.P., I.S., J.D., J.R.; Investigation: S.P., I.S., S.S., M.G.; Visualization: S.P., I.S.; Supervision: I.S.; Writing—original draft: S.P., I.S.; Writing— review & editing: S.P., J.D., S.S., M.G., J.R., I.S.

## Data, code and materials availability

All data and code needed to evaluate and reproduce the results in the paper are present in the Supplementary Materials. This study did not generate any new materials.

## REFERENCES

1. A. T. Nurden, P. Nurden, Congenital platelet disorders and understanding of platelet function. Br J Haematol 165, 165–178 (2014).

2. W. H. Kahr, J. Hinckley, L. Li, H. Schwertz, H. Christensen, J. W. Rowley, F. G. Pluthero, D. Urban, S. Fabbro, B. Nixon, R. Gadzinski, M. Storck, K. Wang, G. Y. Ryu, S. M. Jobe, B. C. Schutte, J. Moseley, N. B. Loughran, J. Parkinson, A. S. Weyrich, J. Di Paola, Mutations in NBEAL2, encoding a BEACH protein, cause gray platelet syndrome. Nat Genet 43, 738–740 (2011).

3. I. Schoen, M. Kenny, S. Patil, Platelet mechanosensing as key to understanding platelet function. Current Opinion in Hematology 31, 24–31 (2024).

4. D. M. Coenen, H. R. Alfar, S. Mohammadmoradi, S. W. Whiteheart, Platelet alpha-granule cargo packaging and endocytosis are important for normal mouse skin wound healing. Blood Adv 10, 2376–2394 (2026).

5. C. Deppermann, D. Cherpokova, P. Nurden, J. N. Schulz, I. Thielmann, P. Kraft, T. Vogtle, C. Kleinschnitz, S. Dutting, G. Krohne, S. A. Eming, A. T. Nurden, B. Eckes, G. Stoll, D. Stegner, B. Nieswandt, Gray platelet syndrome and defective thrombo-inflammation in Nbeal2-deficient mice. J Clin Invest 123, 3331–3342 (2013).

6. L. Nicolai, S. Massberg, Platelets as key players in inflammation and infection. Curr Opin Hematol 27, 34–40 (2020).

7. L. Yue, Z. Pang, H. Li, T. Yang, L. Guo, L. Liu, J. Mei, X. Song, T. Xie, Y. Zhang, X. He, T. J. Lin, Z. Xie, CXCL4 contributes to host defense against acute Pseudomonas aeruginosa lung infection. PloS one 13, e0205521 (2018).

8. V. Martin-Granado, S. Ortiz-Rivero, R. Carmona, S. Gutierrez-Herrero, M. Barrera, L. San-Segundo, C. Sequera, P. Perdiguero, F. Lozano, F. Martin-Herrero, J. R. Gonzalez-Porras, R. Munoz-Chapuli, A. Porras, C. Guerrero, C3G promotes a selective release of angiogenic factors from activated mouse platelets to regulate angiogenesis and tumor metastasis. Oncotarget 8, 110994–111011 (2017).

9. M. Labelle, S. Begum, R. O. Hynes, Direct signaling between platelets and cancer cells induces an epithelial-mesenchymal-like transition and promotes metastasis. Cancer cell 20, 576–590 (2011).

10. A. Eckly, J. Y. Rinckel, F. Proamer, N. Ulas, S. Joshi, S. W. Whiteheart, C. Gachet, Respective contributions of single and compound granule fusion to secretion by activated platelets. Blood 128, 2538–2549 (2016).

11. I. D. Pokrovskaya, S. Joshi, M. Tobin, R. Desai, M. A. Aronova, J. A. Kamykowski, G. Zhang, S. W. Whiteheart, R. D. Leapman, B. Storrie, SNARE-dependent membrane fusion initiates alpha-granule matrix decondensation in mouse platelets. Blood Adv 2, 2947–2958 (2018).

12. D. Jonnalagadda, L. T. Izu, S. W. Whiteheart, Platelet secretion is kinetically heterogeneous in an agonist-responsive manner. Blood 120, 5209–5216 (2012).

13. S. Koseoglu, C. G. Peters, J. L. Fitch-Tewfik, O. Aisiku, L. Danglot, T. Galli, R. Flaumenhaft, VAMP-7 links granule exocytosis to actin reorganization during platelet activation. Blood 126, 651–660 (2015).

14. Y. Sakurai, J. L. Fitch-Tewfik, Y. Qiu, B. Ahn, D. R. Myers, R. Tran, M. E. Fay, L. Ding, P. W. Spearman, A. D. Michelson, R. Flaumenhaft, W. A. Lam, Platelet geometry sensing spatially regulates alpha-granule secretion to enable matrix self-deposition. Blood 126, 531–538 (2015).

15. J. E. Aslan, O. J. McCarty, Rho GTPases in platelet function. J Thromb Haemost 11, 35–46 (2013).

16. D. Pandey, P. Goyal, S. Dwivedi, W. Siess, Unraveling a novel Rac1-mediated signaling pathway that regulates cofilin dephosphorylation and secretion in thrombin-stimulated platelets. Blood 114, 415–424 (2009).

17. H. Akbar, J. Kim, K. Funk, J. A. Cancelas, X. Shang, L. Chen, J. F. Johnson, D. A. Williams, Y. Zheng, Genetic and pharmacologic evidence that Rac1 GTPase is involved in regulation of platelet secretion and aggregation. J Thromb Haemost 5, 1747–1755 (2007).

18. L. Stefanini, Y. Boulaftali, T. D. Ouellette, M. Holinstat, L. Desire, B. Leblond, P. Andre, P. B. Conley, W. Bergmeier, Rap1-Rac1 circuits potentiate platelet activation. Arterioscler Thromb Vasc Biol 32, 434–441 (2012).

19. C. Leon, A. Eckly, B. Hechler, B. Aleil, M. Freund, C. Ravanat, M. Jourdain, C. Nonne, J. Weber, R. Tiedt, M. P. Gratacap, S. Severin, J. P. Cazenave, F. Lanza, R. Skoda, C. Gachet, Megakaryocyte-restricted MYH9 inactivation dramatically affects hemostasis while preserving platelet aggregation and secretion. Blood 110, 3183–3191 (2007).

20. G. J. Johnson, L. A. Leis, M. D. Krumwiede, J. G. White, The critical role of myosin IIA in platelet internal contraction. J Thromb Haemost 5, 1516–1529 (2007).

21. M. Kenny, A. Y. Pollitt, S. Patil, D. W. Hiebner, A. Smolenski, N. Lakic, R. Fisher, R. Alsufyani, S. Lickert, V. Vogel, I. Schoen, Contractility defects hinder glycoprotein VI-mediated platelet activation and affect platelet functions beyond clot contraction. Res Pract Thromb Haemost 8, 102322 (2024).

22. A. Sharda, R. Flaumenhaft, The life cycle of platelet granules. F1000Res 7, 236 (2018).

23. G. Eitzen, Actin remodeling to facilitate membrane fusion. Biochimica et biophysica acta 1641, 175–181 (2003).

24. M. Paul, K. Golla, H. Kim, Gelsolin Modulates Platelet Dense Granule Secretion and Hemostasis via the Actin Cytoskeleton. Thromb Haemost 123, 219–230 (2023).

25. R. Flaumenhaft, J. R. Dilks, N. Rozenvayn, R. A. Monahan-Earley, D. Feng, A. M. Dvorak, The actin cytoskeleton differentially regulates platelet alpha-granule and dense-granule secretion. Blood 105, 3879–3887 (2005).

26. K. Woronowicz, J. R. Dilks, N. Rozenvayn, L. Dowal, P. S. Blair, C. G. Peters, L. Woronowicz, R. Flaumenhaft, The platelet actin cytoskeleton associates with SNAREs and participates in alpha-granule secretion. Biochemistry 49, 4533–4542 (2010).

27. K. Golla, M. Paul, T. C. Lengyell, E. M. Simpson, H. Falet, H. Kim, A novel association between platelet filamin A and soluble N-ethylmaleimide sensitive factor attachment proteins regulates granule secretion. Res Pract Thromb Haemost 7, 100019 (2023).

28. C. Fernandez-Infante, L. Hernandez-Cano, O. Herranz, P. Berrocal, C. Sicilia-Navarro, J. R. Gonzalez-Porras, J. M. Bastida, A. Porras, C. Guerrero, Platelet C3G: a key player in vesicle exocytosis, spreading and clot retraction. Cell Mol Life Sci 81, 84 (2024).

29. C. G. Peters, A. D. Michelson, R. Flaumenhaft, Granule exocytosis is required for platelet spreading: differential sorting of alpha-granules expressing VAMP-7. Blood 120, 199–206 (2012).

30. Z. Li, M. K. Delaney, K. A. O’Brien, X. Du, Signaling during platelet adhesion and activation. Arterioscler Thromb Vasc Biol 30, 2341–2349 (2010).

31. M. Nagy, G. Perrella, A. Dalby, M. F. Becerra, L. Garcia Quintanilla, J. A. Pike, N. V. Morgan, E. E. Gardiner, J. W. M. Heemskerk, L. Azocar, J. F. Miquel, D. Mezzano, S. P. Watson, Flow studies on human GPVI-deficient blood under coagulating and noncoagulating conditions. Blood Adv 4, 2953–2961 (2020).

32. J. Cho, D. F. Mosher, Impact of fibronectin assembly on platelet thrombus formation in response to type I collagen and von Willebrand factor. Blood 108, 2229–2236 (2006).

33. S. Lickert, M. Kenny, K. Selcuk, J. L. Mehl, M. Bender, S. M. Fruh, M. A. Burkhardt, J. D. Studt, B. Nieswandt, I. Schoen, V. Vogel, Platelets drive fibronectin fibrillogenesis using integrin alphaIIbbeta3. Sci Adv 8, eabj8331 (2022).

34. A. Aburima, M. Berger, B. E. J. Spurgeon, B. A. Webb, K. S. Wraith, M. Febbraio, A. W. Poole, K. M. Naseem, Thrombospondin-1 promotes hemostasis through modulation of cAMP signaling in blood platelets. Blood 137, 678–689 (2021).

35. K. Kojok, S. E. Akoum, M. Mohsen, W. Mourad, Y. Merhi, CD40L Priming of Platelets via NF-kappaB Activation is CD40- and TAK1-Dependent. J Am Heart Assoc 7, e03677 (2018).

36. F. Selheim, H. Holmsen, F. S. Vassbotn, Identification of functional VEGF receptors on human platelets. FEBS Lett 512, 107–110 (2002).

37. F. S. Vassbotn, O. K. Havnen, C. H. Heldin, H. Holmsen, Negative feedback regulation of human platelets via autocrine activation of the platelet-derived growth factor alpha-receptor. The Journal of biological chemistry 269, 13874–13879 (1994).

38. J. E. Italiano, Jr., J. L. Richardson, S. Patel-Hett, E. Battinelli, A. Zaslavsky, S. Short, S. Ryeom, J. Folkman, G. L. Klement, Angiogenesis is regulated by a novel mechanism: pro- and antiangiogenic proteins are organized into separate platelet alpha granules and differentially released. Blood 111, 1227–1233 (2008).

39. J. Kamykowski, P. Carlton, S. Sehgal, B. Storrie, Quantitative immunofluorescence mapping reveals little functional coclustering of proteins within platelet alpha-granules. Blood 118, 1370–1373 (2011).

40. E. M. Cramer, G. Berger, M. C. Berndt, Platelet alpha-granule and plasma membrane share two new components: CD9 and PECAM-1. Blood 84, 1722–1730 (1994).

41. H. van Nispen tot Pannerden, F. de Haas, W. Geerts, G. Posthuma, S. van Dijk, H. F. Heijnen, The platelet interior revisited: electron tomography reveals tubular alpha-granule subtypes. Blood 116, 1147–1156 (2010).

42. J. Chung, D. Jeong, G. H. Kim, S. Go, J. Song, E. Moon, Y. H. Huh, D. Kim, Super-resolution imaging of platelet-activation process and its quantitative analysis. Sci Rep 11, 10511 (2021).

43. E. L. Faulkner, J. A. Pike, E. Garlick, R. K. Neely, I. B. Styles, S. P. Watson, N. S. Poulter, S. G. Thomas, Expansion microscopy allows quantitative characterization of structural organization of platelet aggregates. J Thromb Haemost 23, 2618–2633 (2025).

44. H. S. Heil, M. Aigner, S. Maier, P. Gupta, L. M. C. Evers, V. Gob, C. Kusch, M. Meub, B. Nieswandt, D. Stegner, K. G. Heinze, Mapping densely packed alphaIIbbeta3 receptors in murine blood platelets with expansion microscopy. Platelets 33, 849–858 (2022).

45. N. S. Poulter, A. Y. Pollitt, A. Davies, D. Malinova, G. B. Nash, M. J. Hannon, Z. Pikramenou, J. Z. Rappoport, J. H. Hartwig, D. M. Owen, A. J. Thrasher, S. P. Watson, S. G. Thomas, Platelet actin nodules are podosome-like structures dependent on Wiskott-Aldrich syndrome protein and ARP2/3 complex. Nature communications 6, 7254 (2015).

46. S. Lickert, S. Sorrentino, J. D. Studt, O. Medalia, V. Vogel, I. Schoen, Morphometric analysis of spread platelets identifies integrin alpha(IIb)beta(3)-specific contractile phenotype. Sci Rep 8, 5428 (2018).

47. S. Go, D. Jeong, J. Chung, G. H. Kim, J. Song, E. Moon, Y. H. Huh, D. Kim, Super-resolution imaging reveals cytoskeleton-dependent organelle rearrangement within platelets at intermediate stages of maturation. Structure 29, 810–822 e813 (2021).

48. D. Westmoreland, M. Shaw, W. Grimes, D. J. Metcalf, J. J. Burden, K. Gomez, A. E. Knight, D. F. Cutler, Super-resolution microscopy as a potential approach to diagnosis of platelet granule disorders. J Thromb Haemost 14, 839–849 (2016).

49. H. Chanzu, J. Lykins, S. Wigna-Kumar, S. Joshi, I. Pokrovskaya, B. Storrie, G. Pejler, J. P. Wood, S. W. Whiteheart, Platelet alpha-granule cargo packaging and release are affected by the luminal proteoglycan, serglycin. J Thromb Haemost 19, 1082–1095 (2021).

50. F. G. Pluthero, W. H. A. Kahr, Evaluation of human platelet granules by structured illumination laser fluorescence microscopy. Platelets 34, 2157808 (2023).

51. M. Swinkels, F. Atiq, P. E. Burgisser, J. A. Slotman, A. B. Houtsmuller, C. de Heus, J. Klumperman, F. W. G. Leebeek, J. Voorberg, A. J. G. Jansen, R. Bierings, Quantitative 3D microscopy highlights altered von Willebrand factor alpha-granule storage in patients with von Willebrand disease with distinct pathogenic mechanisms. Res Pract Thromb Haemost 5, e12595 (2021).

52. M. Swinkels, S. Hordijk, P. E. Burgisser, J. A. Slotman, T. Carter, F. W. G. Leebeek, A. J. G. Jansen, J. Voorberg, R. Bierings, Quantitative super-resolution imaging of platelet degranulation reveals differential release of von Willebrand factor and von Willebrand factor propeptide from alpha-granules. J Thromb Haemost 21, 1967–1980 (2023).

53. N. S. Poulter, A. Y. Pollitt, D. M. Owen, E. E. Gardiner, R. K. Andrews, H. Shimizu, D. Ishikawa, D. Bihan, R. W. Farndale, M. Moroi, S. P. Watson, S. M. Jung, Clustering of glycoprotein VI (GPVI) dimers upon adhesion to collagen as a mechanism to regulate GPVI signaling in platelets. J Thromb Haemost 15, 549–564 (2017).

54. J. Bergstrand, L. Xu, X. Miao, N. Li, O. Oktem, B. Franzen, G. Auer, M. Lomnytska, J. Widengren, Super-resolution microscopy can identify specific protein distribution patterns in platelets incubated with cancer cells. Nanoscale 11, 10023–10033 (2019).

55. T. M. Svitkina, E. A. Bulanova, O. Y. Chaga, D. M. Vignjevic, S. Kojima, J. M. Vasiliev, G. G. Borisy, Mechanism of filopodia initiation by reorganization of a dendritic network. J Cell Biol 160, 409–421 (2003).

56. A. Obergfell, B. A. Judd, M. A. del Pozo, M. A. Schwartz, G. A. Koretzky, S. J. Shattil, The molecular adapter SLP-76 relays signals from platelet integrin alphaIIbbeta3 to the actin cytoskeleton. The Journal of biological chemistry 276, 5916–5923 (2001).

57. Y. Qiu, A. C. Brown, D. R. Myers, Y. Sakurai, R. G. Mannino, R. Tran, B. Ahn, E. T. Hardy, M. F. Kee, S. Kumar, G. Bao, T. H. Barker, W. A. Lam, Platelet mechanosensing of substrate stiffness during clot formation mediates adhesion, spreading, and activation. Proceedings of the National Academy of Sciences of the United States of America 111, 14430–14435 (2014).

58. D. R. Myers, Y. Qiu, M. E. Fay, M. Tennenbaum, D. Chester, J. Cuadrado, Y. Sakurai, J. Baek, R. Tran, J. C. Ciciliano, B. Ahn, R. G. Mannino, S. T. Bunting, C. Bennett, M. Briones, A. Fernandez-Nieves, M. L. Smith, A. C. Brown, T. Sulchek, W. A. Lam, Single-platelet nanomechanics measured by high-throughput cytometry. Nature materials 16, 230–235 (2017).

59. O. Inoue, K. Suzuki-Inoue, W. L. Dean, J. Frampton, S. P. Watson, Integrin alpha2beta1 mediates outside-in regulation of platelet spreading on collagen through activation of Src kinases and PLCgamma2. J Cell Biol 160, 769–780 (2003).

60. B. Jimenez, O. V. Volpert, S. E. Crawford, M. Febbraio, R. L. Silverstein, N. Bouck, Signals leading to apoptosis-dependent inhibition of neovascularization by thrombospondin-1. Nat Med 6, 41–48 (2000).

61. Z. Li, L. He, K. Wilson, D. Roberts, Thrombospondin-1 inhibits TCR-mediated T lymphocyte early activation. J Immunol 166, 2427–2436 (2001).

62. N. J. Galvin, P. M. Vance, V. M. Dixit, B. Fink, W. A. Frazier, Interaction of human thrombospondin with types I-V collagen: direct binding and electron microscopy. J Cell Biol 104, 1413–1422 (1987).

63. S. Rosini, N. Pugh, A. M. Bonna, D. J. S. Hulmes, R. W. Farndale, J. C. Adams, Thrombospondin-1 promotes matrix homeostasis by interacting with collagen and lysyl oxidase precursors and collagen cross-linking sites. Sci Signal 11, (2018).

64. W. Roberts, S. Magwenzi, A. Aburima, K. M. Naseem, Thrombospondin-1 induces platelet activation through CD36-dependent inhibition of the cAMP/protein kinase A signaling cascade. Blood 116, 4297–4306 (2010).

65. M. Lopes da Silva, D. F. Cutler, von Willebrand factor multimerization and the polarity of secretory pathways in endothelial cells. Blood 128, 277–285 (2016).

66. S. Verhenne, F. Denorme, S. Libbrecht, A. Vandenbulcke, I. Pareyn, H. Deckmyn, A. Lambrecht, B. Nieswandt, C. Kleinschnitz, K. Vanhoorelbeke, S. F. De Meyer, Platelet-derived VWF is not essential for normal thrombosis and hemostasis but fosters ischemic stroke injury in mice. Blood 126, 1715–1722 (2015).

67. N. Dhanesha, P. Prakash, P. Doddapattar, I. Khanna, M. J. Pollpeter, M. K. Nayak, J. M. Staber, A. K. Chauhan, Endothelial Cell-Derived von Willebrand Factor Is the Major Determinant That Mediates von Willebrand Factor-Dependent Acute Ischemic Stroke by Promoting Postischemic Thrombo-Inflammation. Arterioscler Thromb Vasc Biol 36, 1829–1837 (2016).

68. D. W. Hiebner, S. Patil, M. Kenny, J. Baumann, F. Mullin, S. Benn, S. Lickert, V. Vogel, M. Bender, I. Schoen, Regulation of platelet contractility by agonists present across a thrombus. Blood Advances 10, 3442–3456 (2026).

69. I. D. Pokrovskaya, S. Yadav, A. Rao, E. McBride, J. A. Kamykowski, G. Zhang, M. A. Aronova, R. D. Leapman, B. Storrie, 3D ultrastructural analysis of alpha-granule, dense granule, mitochondria, and canalicular system arrangement in resting human platelets. Res Pract Thromb Haemost 4, 72–85 (2020).

70. J. A. Berriman, S. Li, L. J. Hewlett, S. Wasilewski, F. N. Kiskin, T. Carter, M. J. Hannah, P. B. Rosenthal, Structural organization of Weibel-Palade bodies revealed by cryo-EM of vitrified endothelial cells. Proceedings of the National Academy of Sciences of the United States of America 106, 17407–17412 (2009).

71. C. Yeaman, K. K. Grindstaff, W. J. Nelson, Mechanism of recruiting Sec6/8 (exocyst) complex to the apical junctional complex during polarization of epithelial cells. Journal of cell science 117, 559–570 (2004).

72. M. Kawato, R. Shirakawa, H. Kondo, T. Higashi, T. Ikeda, K. Okawa, S. Fukai, O. Nureki, T. Kita, H. Horiuchi, Regulation of platelet dense granule secretion by the Ral GTPase-exocyst pathway. The Journal of biological chemistry 283, 166–174 (2008).

73. K. K. Grindstaff, C. Yeaman, N. Anandasabapathy, S. C. Hsu, E. Rodriguez-Boulan, R. H. Scheller, W. J. Nelson, Sec6/8 complex is recruited to cell-cell contacts and specifies transport vesicle delivery to the basal-lateral membrane in epithelial cells. Cell 93, 731–740 (1998).

74. M. Rodrigues, N. Kosaric, C. A. Bonham, G. C. Gurtner, Wound Healing: A Cellular Perspective. Physiol Rev 99, 665–706 (2019).

75. F. H. Furby, M. C. Berndt, P. A. Castaldi, J. Koutts, Characterization of calcium-dependent binding of endogenous factor VIII/von Willebrand factor to surface activated platelets. Thrombosis Research 35, 501–511 (1984).

76. H. R. Gralnick, S. B. Williams, L. P. McKeown, L. Magruder, K. Hansmann, M. Vail, R. I. Parker, Platelet von Willebrand Factor. Mayo Clinic Proceedings 66, 634–640 (1991).

77. P. Blair, R. Flaumenhaft, Platelet alpha-granules: basic biology and clinical correlates. Blood Rev 23, 177–189 (2009).

78. H. Fu, Y. Jiang, D. Yang, F. Scheiflinger, W. P. Wong, T. A. Springer, Flow-induced elongation of von Willebrand factor precedes tension-dependent activation. Nature communications 8, 324 (2017).

79. S. F. De Meyer, N. Vandeputte, I. Pareyn, I. Petrus, P. J. Lenting, M. K. Chuah, T. VandenDriessche, H. Deckmyn, K. Vanhoorelbeke, Restoration of plasma von Willebrand factor deficiency is sufficient to correct thrombus formation after gene therapy for severe von Willebrand disease. Arterioscler Thromb Vasc Biol 28, 1621–1626 (2008).

80. E. J. Bowie, L. A. Solberg, Jr., D. N. Fass, C. M. Johnson, G. J. Knutson, M. L. Stewart, L. J. Zoecklein, Transplantation of normal bone marrow into a pig with severe von Willebrand’s disease. J Clin Invest 78, 26–30 (1986).

81. E. Fressinaud, D. Baruch, C. Rothschild, H. Baumgartner, D. Meyer, Platelet von Willebrand factor: evidence for its involvement in platelet adhesion to collagen. Blood 70, 1214–1217 (1987).

82. A. Laroche, N. Rovito, A. Liu, I. Allaeys, L. Heitsch, I. Portier, O. Adeoye, S. R. Pizzella, J. A. Giles, J. Di Paola, J. M. Lee, D. Bark, R. A. Campbell, D. Mukherjee, E. Boilard, F. Denorme, Mechanisms of von Willebrand factor activation driving no reflow in ischemic stroke. Proceedings of the National Academy of Sciences of the United States of America 123, e2610397123 (2026).

83. R. T. McGrath, E. McRae, O. P. Smith, J. S. O’Donnell, Platelet von Willebrand factor – structure, function and biological importance. British Journal of Haematology 148, 834–843 (2010).

84. J. A. Dent, M. Galbusera, Z. M. Ruggeri, Heterogeneity of plasma von Willebrand factor multimers resulting from proteolysis of the constituent subunit. The Journal of Clinical Investigation 88, 774–782 (1991).

85. N. A. Zainal Abidin, E. K. W. Poon, C. Szydzik, M. Timofeeva, F. Akbaridoust, R. J. Brazilek, F. J. Tovar Lopez, X. Ma, C. Lav, I. Marusic, P. E. Thompson, A. Mitchell, A. S. H. Ooi, J. R. Hamilton, W. S. Nesbitt, An extensional strain sensing mechanosome drives adhesion-independent platelet activation at supraphysiological hemodynamic gradients. BMC Biology 20, 73 (2022).

86. M. Tomaiuolo, T. J. Stalker, J. D. Welsh, S. L. Diamond, T. Sinno, L. F. Brass, A systems approach to hemostasis: 2. Computational analysis of molecular transport in the thrombus microenvironment. Blood 124, 1816–1823 (2014).

87. R. Sandmann, S. Koster, Topographic Cues Reveal Two Distinct Spreading Mechanisms in Blood Platelets. Sci Rep 6, 22357 (2016).

88. R. D. Allen, L. R. Zacharski, S. T. Widirstky, R. Rosenstein, L. M. Zaitlin, D. R. Burgess, Transformation and motility of human platelets: details of the shape change and release reaction observed by optical and electron microscopy. J Cell Biol 83, 126–142 (1979).

89. K. Croce, R. Flaumenhaft, M. Rivers, B. Furie, B. C. Furie, I. M. Herman, D. A. Potter, Inhibition of Calpain Blocks Platelet Secretion, Aggregation, and Spreading *. Journal of Biological Chemistry 274, 36321–36327 (1999).

90. I. Pleines, A. Eckly, M. Elvers, I. Hagedorn, S. Eliautou, M. Bender, X. Wu, F. Lanza, C. Gachet, C. Brakebusch, B. Nieswandt, Multiple alterations of platelet functions dominated by increased secretion in mice lacking Cdc42 in platelets. Blood 115, 3364–3373 (2010).

91. H. Akbar, X. Shang, R. Perveen, M. Berryman, K. Funk, J. F. Johnson, N. N. Tandon, Y. Zheng, Gene targeting implicates Cdc42 GTPase in GPVI and non-GPVI mediated platelet filopodia formation, secretion and aggregation. PloS one 6, e22117 (2011).

92. M. Machacek, L. Hodgson, C. Welch, H. Elliott, O. Pertz, P. Nalbant, A. Abell, G. L. Johnson, K. M. Hahn, G. Danuser, Coordination of Rho GTPase activities during cell protrusion. Nature 461, 99–103 (2009).

93. A. E. Urban, E. O. Quick, K. P. Miller, J. Krcmery, H.-G. Simon, Pdlim7 Regulates Arf6-Dependent Actin Dynamics and Is Required for Platelet-Mediated Thrombosis in Mice. PloS one 11, e0164042 (2016).

94. R. Crombie, R. L. Silverstein, C. MacLow, S. F. A. Pearce, R. L. Nachman, J. Laurence, Identification of a CD36-related Thrombospondin 1–binding Domain in HIV-1 Envelope Glycoprotein gp120: Relationship to HIV-1–specific Inhibitory Factors in Human Saliva. Journal of Experimental Medicine 187, 25–35 (1998).

95. M. Kenny, S. Stamboroski, R. Taher, D. Bruggemann, I. Schoen, Nanofiber Topographies Enhance Platelet-Fibrinogen Scaffold Interactions. Advanced healthcare materials 11, e2200249 (2022).

96. R. M. Power, A. Tschanz, T. Zimmermann, J. Ries, Build and operation of a custom 3D, multicolor, single-molecule localization microscope. Nat Protoc 19, 2467–2525 (2024).

97. J. Ries, SMAP: a modular super-resolution microscopy analysis platform for SMLM data. Nat Methods 17, 870–872 (2020).

98. Y. Li, M. Mund, P. Hoess, J. Deschamps, U. Matti, B. Nijmeijer, V. J. Sabinina, J. Ellenberg, I. Schoen, J. Ries, Real-time 3D single-molecule localization using experimental point spread functions. Nature Methods 15, 367–369 (2018).

