## Supplementary Figures S1-S8 for "Super-resolution mapping of platelet α-granules uncovers non-polarized secretion and differential retention of VWF and thrombospondin-1"

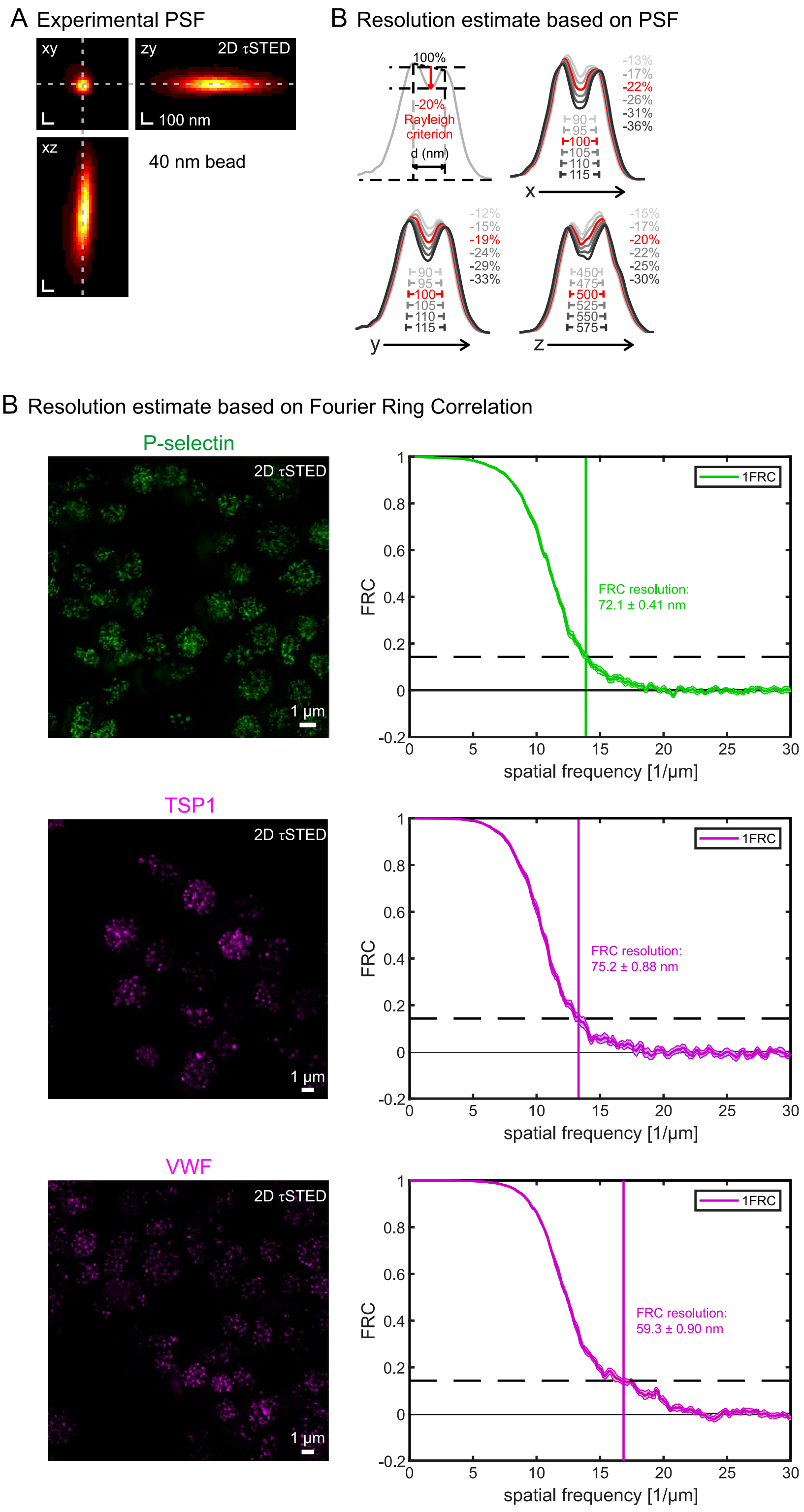


**Supplementary Figure S1**. Resolution estimation for 2D τSTED images of α-granule stainings in resting platelets.

**A**. Experimental point spread function (PSF). Z-stacks of 40 nm Abberior Star635P beads were acquired using standard 2D τSTED settings. z-stacks of 6 independent beads were used to construct an average spline-interpolated PSF. Shown are orthogonal views of this average PSF. Please note that the experimentally measured PSF is a convolution of the true PSF with the size of the fluorescent beads.

**B**. Resolution estimate based on the experimental PSF. Two point sources at distance *d* (in different directions) were convolved using the experimental PSF. Shown are central intensity line profiles along the direction. Rayleigh's criterion, namely a minimal drop of the intensity by -20% between the intensity maxima, was used to estimate the resolution. As expected, the 2D STED modality mainly increases the lateral resolution by more than 2-fold but only slightly increases the axial resolution compared to confocal imaging. Please note that the lateral resolution estimate derived from the experimentally measured PSF is an underestimate of the true resolution due to the convolution with the finite fluorescent bead size.

**C**. Lateral resolution estimates directly derived from experimental images using the single-image Fourier Ring Correlation (1FRC) method. P-selectin (Psel), thrombospondin-1 (TSP1) or von Willebrand Factor (VWF) were stained with Abberior Star635P. Images were acquired using standard 2D $\tau$STED settings. FRC analysis shows a routine lateral resolution better than 75 nm, depending on signal strength. This resolution estimate is in excellent agreement with the PSF-based estimate when considering the bead size (40 nm).

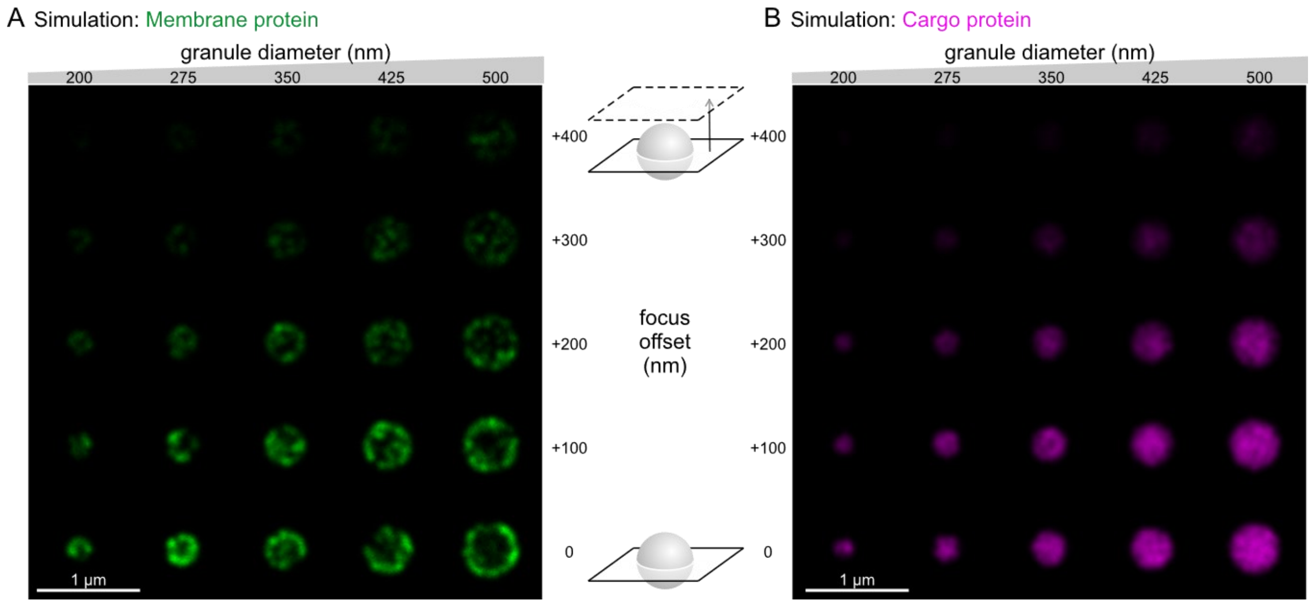


**Supplementary Figure S2**. Effect of focus offset on granule appearance in simulated 2D τSTED images. Spheres ranged from 200-500 nm in diameter, the focus offset from 0-400 nm.

**A**. Simulated images of protein markers residing on the membrane of granules.

**B**. Simulated images of cargo proteins filling the inside of granules.

Due to the elongated PSF in z, the 3-dimensional structure of granules remains ill-resolved, and the main consequence of defocusing is a reduction in signal intensity.

Correction for bleedthrough

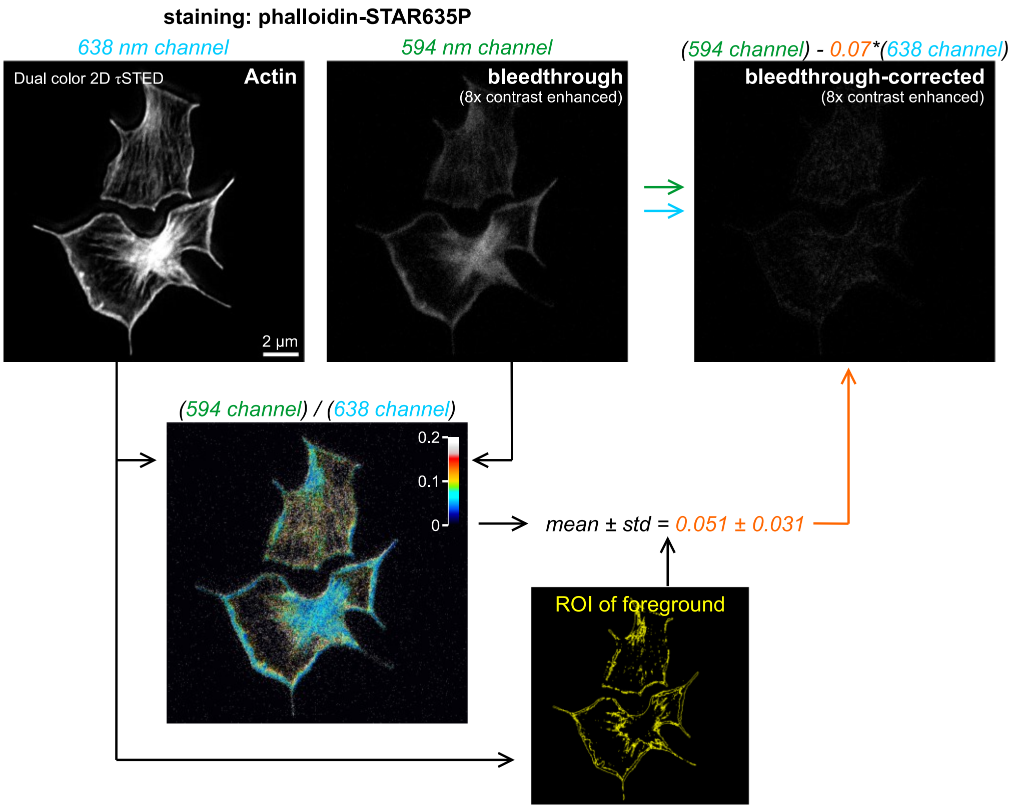


**Supplementary Figure S3**. Post-acquisition correction of channel crosstalk in 2C τSTED images. Due to spectral overlap of STAR635P with the 594 nm detection channel (used to image AF594), and some direct excitation of STAR635P by the 494 nm excitation line, the STAR635P staining is also weakly detected in the 594 nm channel. There is no crosstalk of the AF594 stain into the 638 nm channel since the dye is not excited by the 638 nm excitation line. To correct for this crosstalk, we take advantage of the constant acquisition settings used across all experiments (594/638 nm excitation powers, spectral detection windows, detector settings, depletion laser power). A sample was exclusively stained with phalloidin-STAR635P (no AF594 staining) and imaged in both channels. The 594 nm channel image was divided pixel-by-pixel by the 638 nm channel image. A region of interest (ROI) was defined based on the 638 nm signal to limit the measurements of the intensity ratio (594/638) to the stained subregion. The derived correction factor was used to correct the 594 nm channel image from the crosstalk by subtracting the rescaled 638 nm channel image. A correction factor of 0.07 was used, which was a conservative choice to remove all possible artifacts.

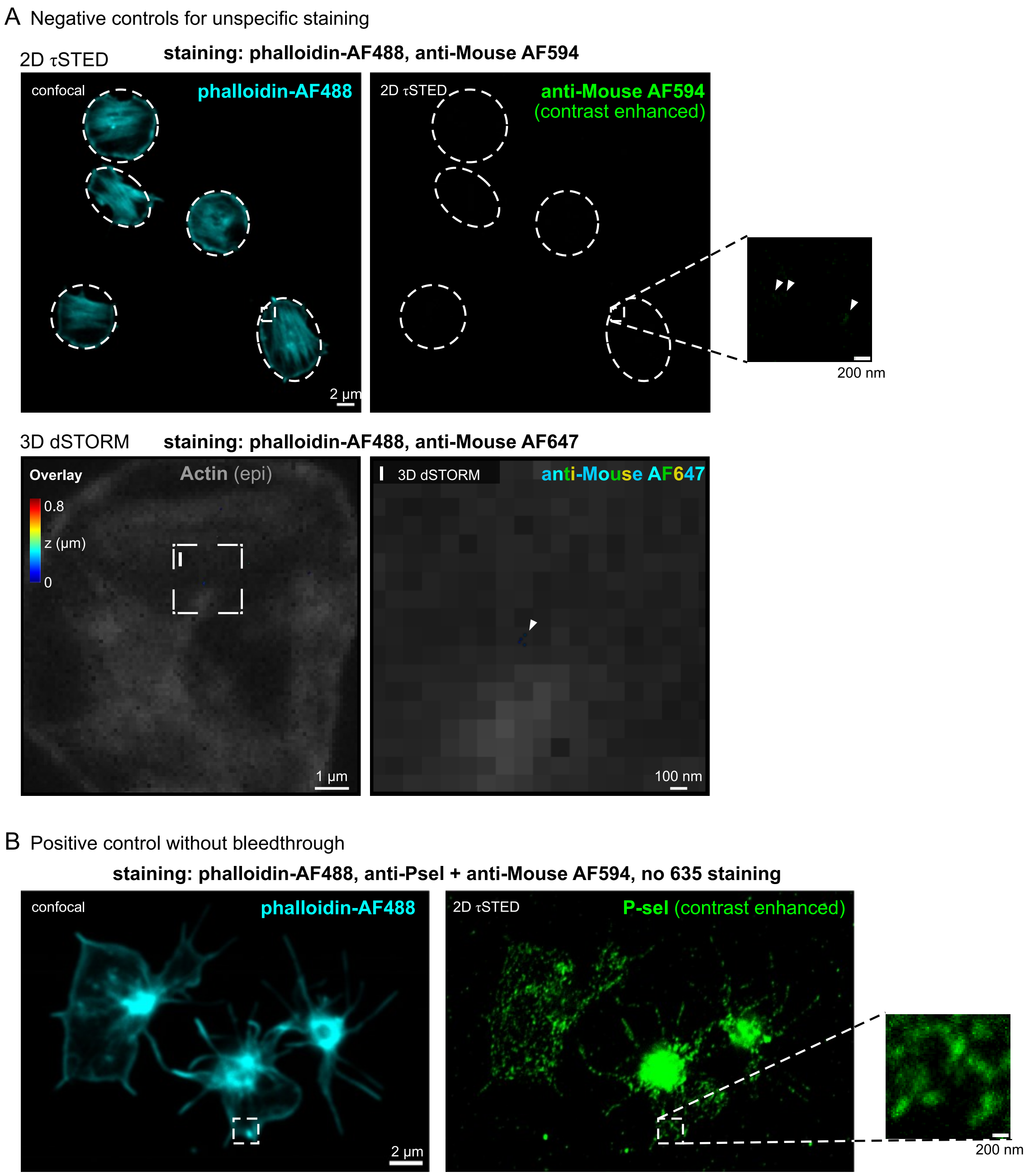


**Supplementary Figure S4**. Experimental controls for superresolution microscopy techniques.
**A**. Negative controls for unspecific staining. Fixed spread platelets were treated the same way as for normal stainings with the exception of not adding a primary antibody. Very few, faint, non-specific signals were observed (arrowheads) which did not interfere with the analysis/interpretation of the specific staining in other experiments.
**B**. Positive control for disperse P-selectin signals. A sample was only stained with AF594 for P-selectin, and single-colour τSTED was performed. Disperse spotty P-selectin signals along filopodia and in lamellipodia were clearly resolved. These signals were brighter and much more frequent than unspecific signals (see subfigure A above), indicating specific staining originating from individual or few P-selectin molecules. The close agreement between this control and the 2C τSTED images rule out that the singals in 2C τSTED images originated from an insufficient bleedthrough correction.

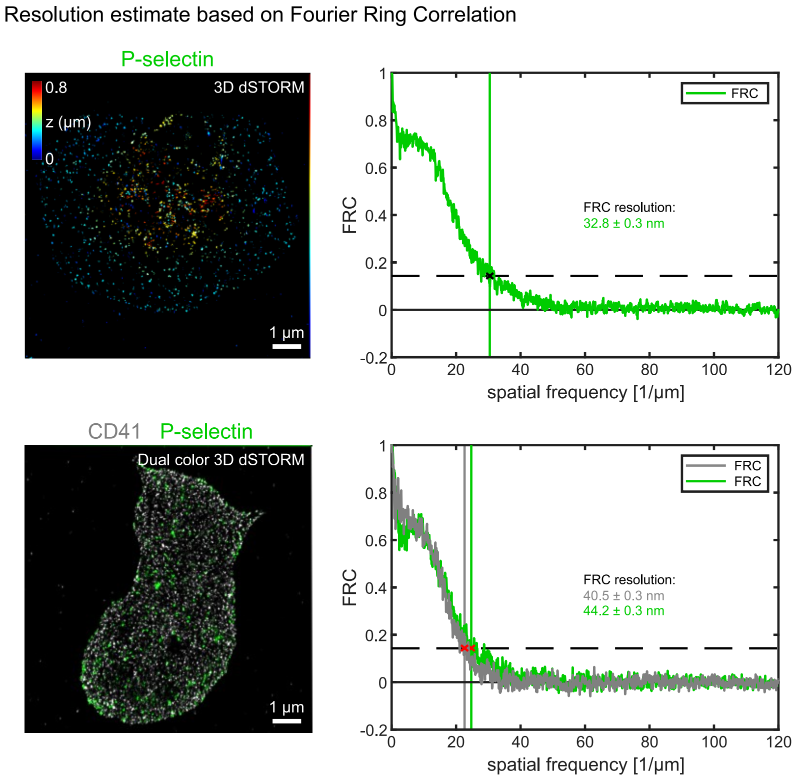


**Supplementary Figure S5**. Lateral resolution estimation for 3D dSTORM images.

Lateral resolution estimates were directly derived from localization data of single color 3D dSTORM or dual color 3D dSTORM measurements using the Fourier Ring Correlation (FRC) method. Stainings were for P-selectin (green) or CD41 (gray). Images were acquired using standard 3D dSTORM settings. FRC analysis shows a routine lateral resolution better than 35 nm in single color and better than 45 nm in dual color measurements.

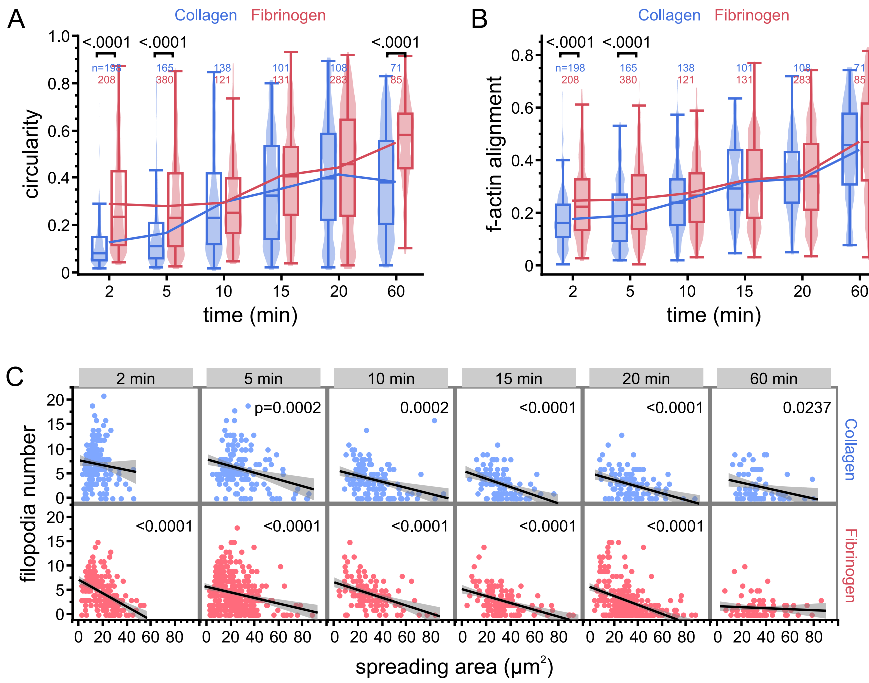


**Supplementary Figure S6**. Comparison of morphometric features in platelets spreading on coverslips coated with collagen (blue) or fibrinogen (red). Shown are results of morphological single cell measurements from 3 independent experiments; the number of total cells is indicated. Experimental settings were as in Figure 5B.

**A**. Cell circularity.

**B**. F-actin alignment.

The lower circularity and lower f-actin alignment on collagen at early time points reflects the higher number of filopodia in these cells (see Figure 5B).

**C**. Correlation analysis of filopodia number versus spreading area. Each data point represents a single cell. Linear regression (black line) and 95% confidence intervals (gray) show an anti-correlation for all time points and on either adhesion protein coating, with less filopodia in platelets that showed more complete spreading.

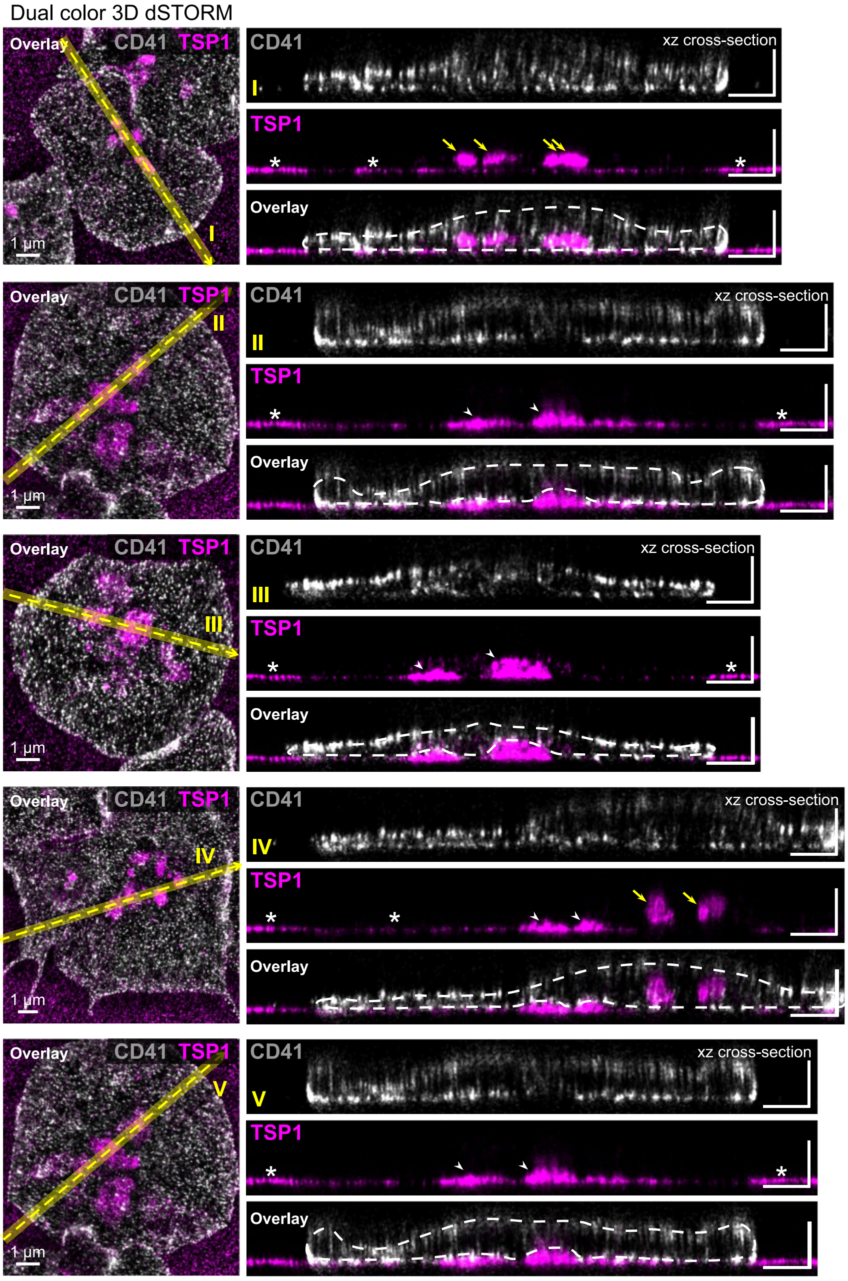


**Supplementary Figure S7**. Basal secretion of TSP1 leads to its retention underneath spread platelets. Additional examples (see also Figure 6B) of dual color 3D STORM measurements of TSP1 (magenta) and CD41 (gray). TSP is detected within the cell in intact granules (yellow arrows), underneath the cell in pockets sealed off by the cell membrane (arrowheads), and adsorbed to the fibrinogen-coated coverslip around platelets and beneath some lamellipodia (asterisks). Without the sideview and the high z-resolution, the ability to distinguish between TSP1 in intact granules and beneath the cell is very limited.

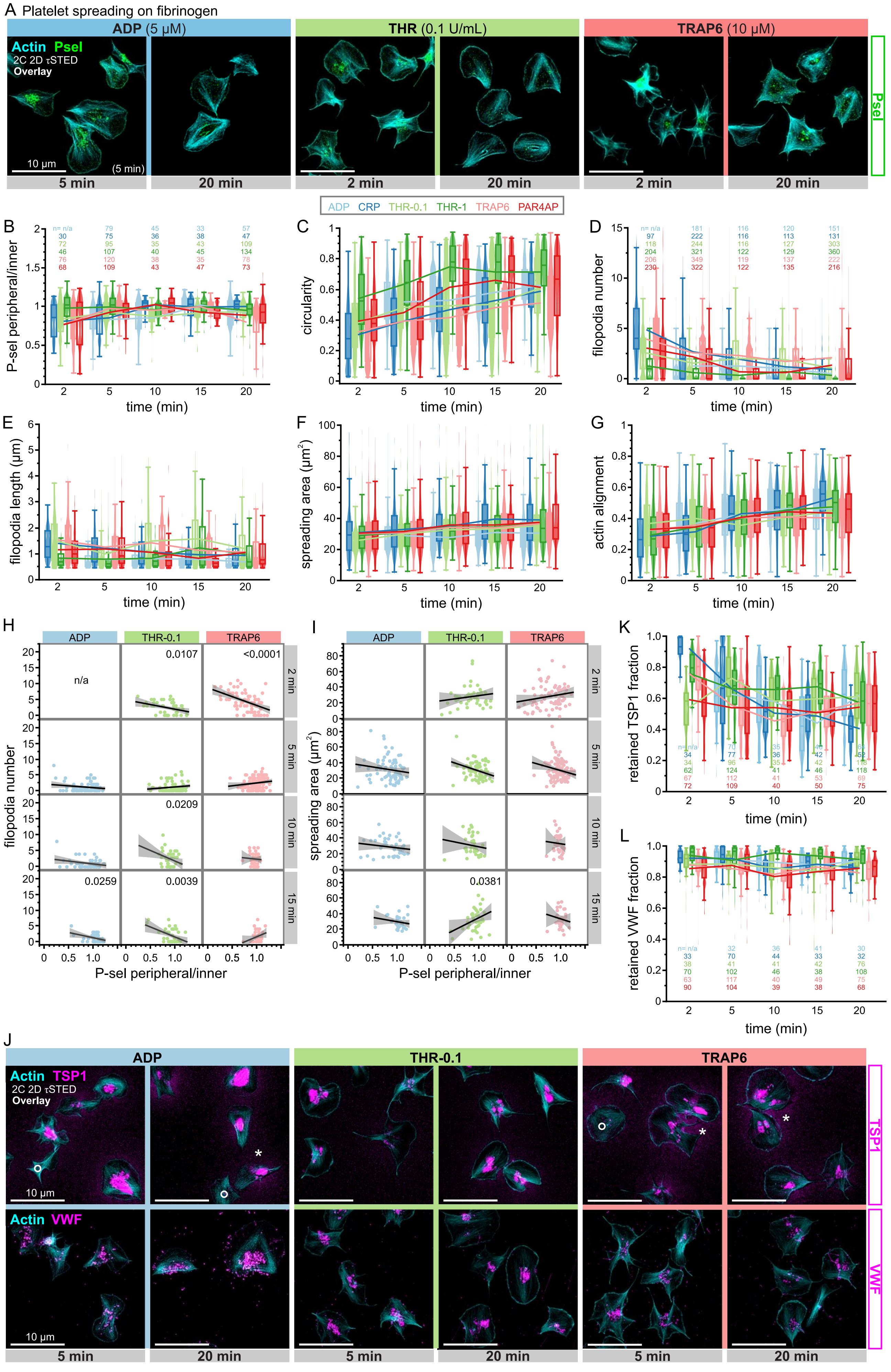


**Supplementary Figure S8.**

**Supplementary Figure S8**. Influence of agonist stimulation during platelet spreading on the coordination of α-granule secretion and cytoskeletal rearrangements: Additional agonists and time points.
Washed human platelets were spread in the presence of agonists on fibrinogen-coated coverslips up to 20 min, fixed at different time points, permeabilized, immunostained, and imaged by SRM.
**A**. Representative SRM images of f-actin (cyan) and P-selectin (green) in human platelets spread on fibrinogen for few (2 or 5) minutes or 20 minutes in the presence of 5 µM ADP (light blue; left), 0.1 U/mL thrombin (light green; middle) or 10 µM TRAP6 (light red; right). Scale bars: 10 µm.
**B**-**G**. Morphometrics of platelets spread in the presence of different agonists (legend) for 2-20 minutes. Single-cell parameters are: **B**. Peripheral-to-central P-selectin intensity ratio; **C**. Circularity; **D**. Filopodia number; **E**. Filopodia length; **F**. spreading area; **G**. Actin alignment. Single cell data are shown as violin plots and summarised as box plots (25/50/75 percentile, whiskers excluding outliers). Connecting lines depict the mean per time point. The total number of platelets per time point pooled from 1-4 independent experiments is denoted above the graphs in (B) for panels (B,H,I) and in (D) for panels (C-G). Statistical tests by ANOVA with Tukey for multiple comparisons. THR-1: thrombin at 1 U/mL; THR-0.1: thrombin at 0.1 U/mL; PAR4AP: PAR4-activating peptide.
**H**-**I**. Correlation analyses of **H**. filopodia number or **I**. spreading area versus the degree of α-granule secretion (measured by the peripheral-to-central P-selectin intensity ratio). An anti-correlation of the filopodia number is observed at 2 minutes, while no significant correlations were seen for the spreading area.
**J**. Representative τSTED images of f-actin (cyan) and thrombospondin 1 (magenta, top row) or VWF (magenta, bottom row) in human platelets spread on fibrinogen for few 5 minutes or 20 minutes in the presence of 5 µM ADP (light blue; left), 0.1 U/mL thrombin (light green; middle) or 10 µM TRAP6 (light red; right). Scale bars: 10 µm.
**K**-**L**. Fraction of **K**. thrombospondin 1 or **L**. VWF retained at the cell compared to its immediate surrounding (for definition, see Figure 6C) for platelets spread in the presence of different agonists. Single cell data are shown as violin plots and summarised as box plots (25/50/75 percentile, whiskers excluding outliers). Connecting lines depict the mean per time point. The total number of platelets per time point is denoted below the graphs. Statistical tests by ANOVA with Tukey for multiple comparisons. THR-1: thrombin at 1 U/mL; THR-0.1: thrombin at 0.1 U/mL; PAR4AP: PAR4-activating peptide.
